# Evolutionary dynamics of neutral phenotypes under DNA substitution models

**DOI:** 10.1101/2020.10.26.355438

**Authors:** Shadi Zabad, Alan M Moses

**Affiliations:** Departments of Computer Science, University of Toronto, Canada; Cell & Systems Biology, University of Toronto, Canada; Ecology and Evolutionary Biology, University of Toronto, Canada; Centre for Analysis of Genome Evolution and Function, University of Toronto, Canada

## Abstract

We study the evolution of quantitative molecular traits in the absence of selection. Using Felsenstein’s 1981 DNA substitution model, we predict a linear restoring force on the mean of an additive phenotype. Remarkably, the mean dynamics are independent of the effect sizes and genotype and are similar to those predicted by the widely-used OU model for stabilizing selection and the house-of-cards model for phenotype evolution. We confirm the predictions empirically using additive molecular phenotypes calculated from ancestral reconstructions of putatively unconstrained DNA sequences in primate genomes. We predict and confirm empirically that the dynamics of the variance are more complicated than those predicted by the OU model, and show that our results for the restoring force of mutation hold even for nonadditive phenotypes, such as number of transcription factor binding sites, longest encoded peptide and folding propensity of the encoded peptide. Our results have implications for efforts to infer selection based on quantitative phenotype dynamics as well as to understand long-term trends in evolution of quantitative molecular traits.

## Introduction

With the increasing availability of high-throughput data about molecular and cellular function from many species, a key challenge is detect selection on these quantitative molecular traits [1]–[9]. The models used for selection in the comparative phylogenetic approach have rich theoretical grounding [10], [11] and software packages are available for data analysis under a variety of inference frameworks [12]–[14]. However, despite evidence that molecular phenotypes may evolve with little selective constraint [15]–[17] and classical work in the area[18], relatively little attention has been paid to improving the realism of the “neutral” or “null hypotheses” for phenotype evolution, as compared to protein coding sequences[19], [20] where the null hypothesis can be explicitly formulated based on empirical estimates from neutral sites [19] or, more recently, the biophysics of protein folding (e.g.,[21]).

The widely-adopted Brownian Motion (also referred to as a “random walk”[22]) model is an appealing null hypothesis for phenotype evolution in comparative analysis of quantitative molecular traits[8]. It is mathematically convenient [23] and is referred to as a model of “pure drift” [1], [13] because it matches quantitative trait evolution in theory under genetic drift alone[23]. A key prediction of the Brownian Motion model is that changes from the ancestral phenotype are symmetric, and therefore, on average, mean phenotypes are not expected to change over time[24]. On the other hand, it has long been appreciated that even in the absence of selection, mutation is expected to be a weak directional force on phenotype evolution[22], [25] and this has been observed empirically in at least one recent mutation accumulation study[26].

This view is captured in the so-called house-of-cards model [27], which was meant to capture the idea that the force of random mutation would be expected to break down what had been built up by selection. More recently, it has been suggested that mutation could be a constructive force in evolution, leading to non-adaptive increases in biological complexity[28], [29].

With the increasing interest in studying evolution of molecular and cellular traits over long evolutionary time-scales, several theoretical studies have begun to take a closer look at the expected effects of mutation on the phenotype, finding that the equilibrium phenotype is the result of a trade-off between the directional effects of mutation and selection [30]–[32], and that biased mutation can drive the phenotype away from the optimum when selection is not strong[33]. To our knowledge, these predictions have not yet been compared directly with data, but global gene expression variation in mutation accumulation studies appears more consistent with the house-of-cards model[34]. Perhaps most intriguingly, a recent empirical study that estimated the effects of mutations on gene expression levels for 10 genes found asymmetric distributions (inconsistent with Brownian Motion) and used simulations to predict approximately linear change over time in the absence of selection for most genes [35].

In order to gain insight into the appropriate null hypothesis for use in comparative studies of quantitative molecular phenotypes, we first develop a neutral model of phenotype evolution that could be compared directly to observations. We work out the dynamics of additive phenotypes in the weak-mutation regime[36], the regime of most relevance for molecular evolutionary studies. Under Felsenstein’s 1981 model of DNA evolution[37], we predict that mutation acts as a linear restoring force on the mean phenotype, similar to the OU models currently used for stabilizing selection[10] and to the house-of-cards model for phenotype evolution[22]. Next, we develop a paradigm based on ancestral genome reconstruction to directly compare the predictions of these models to observations of molecular phenotype evolution. We find remarkable agreement with observations of molecular phenotypes computed from reconstructed ancestors of putatively neutral sequences in primate genomes, even when the phenotypes are not strictly additive.

## Results

### Neutral dynamics of the mean of an additive phenotype under a DNA substitution model

Our goal is to develop a neutral model of phenotype evolution with sufficient realism so that it can be directly compared to observations of molecular phenotypes obtained from closely related extant species. We therefore sought to derive the dynamics of the moments of the phenotype distribution starting with a standard model of DNA substitution. Thus, we model phenotype evolution as a consequence of genetic changes due to mutation and neutral fixation alone.

We denote the mean phenotype in a population or species with Z and we assume that it linearly depends on the underlying genotype matrix X. In our model, X is an L x K binary matrix, where L is the number of contributing loci and K is the number of distinct alleles at each locus. The binary coding of the genotype matrix follows from our assumption of a monomorphic haploid population, so that *X_jk_* = 1 if the k-th allele is fixed at the j-the locus and zero otherwise. Next, we use *β_jk_* to represent the effect size of the k-th allele at the j-th locus. With this, we can write the mean phenotype in a population or species as:

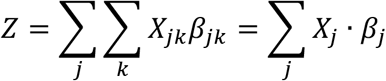

For example, for a DNA-based phenotype with *L* = 10 contributing loci, X and *β* are 10 x 4 matrices. In order to compare with empirical phenotype data, we allow arbitrary values for *β_jk_*, thus making no assumptions about the distribution of effect sizes.

We are interested in examining the dynamics of the mean phenotype as a function of time Z(t). If we assume that the effect sizes remain constant, the time-dependent variable in our model is the underlying genotype matrix X(t):

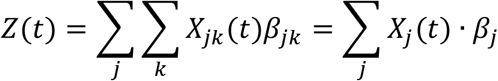

Characterizing the evolution of the genotype matrix in response to mutation, selection, and drift has been a central focus of population genetic theory for the past century[38]. In our model, we make the simplifying assumptions that the only operating evolutionary forces are mutation and drift and that the loci evolve independently according to a continuous-time Markov chain [20] with K states corresponding to the K different alleles. In assuming that a monomorphic initial genotype mutates into another monomorphic genotype we are neglecting the possibility of polymorphism, competition and linkage between genotypes and other population genetic processes. This regime has been referred to as the sequential model [39] and is equivalent to the so-called “weak mutation” assumption commonly used to model molecular evolution at the timescale of inter species divergence [36].

In this work, we use Felsenstein’s 1981 model (F81) of DNA evolution [37] to model the evolution of the individual loci. For each allele k, the model defines the following transition probabilities

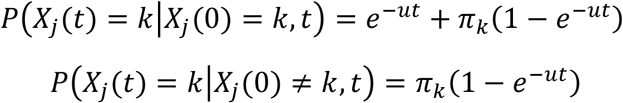

Where *X_j_*(*t*) = *k* denotes that the k-th allele is fixed at the j-th locus at time t and 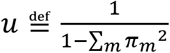 is the DNA substitution rate, scaled so that evolutionary distance, *ut*, is measured in substitutions per site. Given an ancestral population with mean phenotype Z(0), mutation and drift will fix some alleles while driving others to extinction. Assuming that we have an ensemble of replicate populations emanating from that ancestral population, our goal is to provide expressions for the ensemble mean E[Z(t)] and variance, V[Z(t)] of the mean phenotype over time.

In the appendix, we provide detailed derivations that show that the ensemble mean has the following form

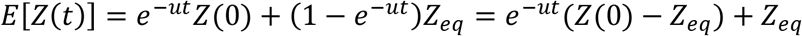

Since we measure evolutionary distance in units of substitutions per site, for evolutionary times of interest in the primate phylogeny considered below, *ut* ≪ 1, so *e^−ut^* ≈ 1 − *ut*. Hence

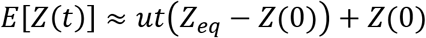

Where 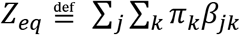 and can be interpreted as the phenotype expected after a long-time of neutral evolution, or the phenotype at “mutational equilibrium.” Thus, under the F81 model, the dynamics of the mean phenotype depend only on the initial phenotype, the phenotype at its mutational equilibrium and the evolutionary distance. Remarkably, we find that the dynamics of the ensemble mean of an additive phenotype is predicted to be *independent* of the starting genotype, X. In other words, the dynamics of the mean phenotype are *the same for all population genotypes* that encode that phenotype. We conjecture that the F81 model is the most realistic DNA substitution model[20] for which this can be true (for example, it is not true of models with transition-transversion rate bias[40], see Discussion). The dynamics of the mean phenotype are also *independent* of the number of alleles, loci, and the distribution of allelic effect sizes, *β*.

Our model predicts that mean phenotype is driven towards the equilibrium with a force simply proportional to the distance of the initial phenotype from its mutational equilibrium, *Z*(0) – *Z_eq_*. Thus, near the mutational equilibrium, mutation can be neglected, but the further the phenotype is from equilibrium, the stronger the directional force of mutation becomes. The appearance of a directional force on the mean phenotype in the absence of selection is contradictory to the Brownian Motion null hypothesis for phenotype evolution[23], which argues that in the absence of selection, the mean phenotype is not expected on average to change in one direction or the other[23]. Importantly, the dynamics of the mean phenotype match exactly the dynamics of the mean of the well-studied Ornstein-Uhlenbeck (OU) process, a stochastic model that is widely used to capture the restoring force of stabilizing selection in phylogenetic comparative analysis[10], [11], [13]. If our prediction of OU-like mean dynamics in the absence of selection is correct, naïve use of the OU process to model selection may be misleading (see discussion).

In the appendix, we provide detailed derivations that show that the ensemble variance of the mean phenotype has the following form:

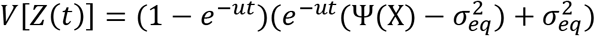

Once again, for evolutionary times of interest in the primate phylogeny considered below, *ut* ≪ 1, so *e^−ut^* ≈ 1 – *ut*. We have

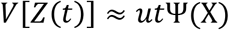

In these expressions, 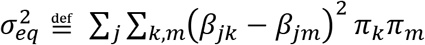, which is the ensemble variance at equilibrium, and 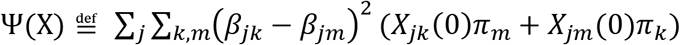 is a new, genotype-dependent quantity whose meaning is not entirely clear, but it captures something about the dispersion of the evolutionary process (see Appendix). In the sums, *k, m* index the unique pairs of alleles. The expressions for the variance make immediately clear that at long times the variance reaches an equilibrium value 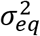, and at short times it increases proportional to evolutionary distance. Importantly, unlike the mean phenotype, whose dynamics depend only on phenotypic quantities, in general, the variance of the mean phenotype *does* depend on the initial genotype through the quantity Ψ(X). In other words, populations with initially identical mean phenotypes, but different genotypes, are predicted to show *identical mean* phenotype evolution, but will potentially show *different variance* evolution. Furthermore, the variance does not always increase monotonically to its equilibrium value. These predictions differ qualitatively from both the Brownian Motion and OU process models typically used in comparative phylogenetic analysis.

Under the assumptions that the genotypes in the population are initially at mutational equilibrium (not likely to be the case for real populations) or that the phenotype is encoded by bi-allelic loci with uniform effect sizes and evolves with symmetric mutation rates (see Appendix) we find that 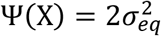 and 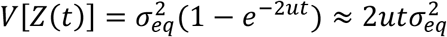 (with the approximation being for *ut* ≪ 1 as above.) Thus, under these assumptions, we do obtain genotype-independent dynamics, whose short-time behavior agrees with the predictions of the Brownian-Motion model [41].

Although in general we predict some differences in the variance dynamics between the F81 and the OU models, we wondered under what circumstances the OU process might provide a good approximation to the variance dynamics of the F81 model. Under the OU model, the variance is

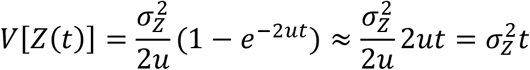

To figure out the fluctuation size, 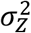, we note that the long-time variance should go to 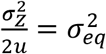. Therefore, under an OU model with only the force of mutation, 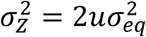 and

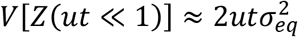

Thus, the OU model (with mutation alone) also matches the predictions of the F81 model exactly under the assumptions of initial genetic equilibrium, or symmetric biallelic mutation with constant effect size. We note, however, that in the F81 model, the force of mutation, u, is never separate from the time, t, implying that these cannot be inferred separately[37]. In the OU model, these appear separately, which could in principle lead additional complexity in inference.

### Molecular phenotypes from ERVs in primate genomes

Since our goal is to test empirically which neutral model should be used in practice, we considered phenotypes that could be computed directly from DNA sequences: these represent so-called quantitative molecular traits[5] with known genetic architecture and no measurement noise. This experimental set-up is attractive for several reasons: we can be sure that the phenotypes are truly “additive” (therefore not violating the assumptions of the model), we can rule out that stochastic effects we observe are intrinsic to the evolutionary process and not due to measurement or sampling errors[42], [43], and we can compare the observations from real DNA sequences to comparable observations from forward simulations under the DNA substitution models to determine the effects (if any) of mis-specification of the substitution model.

To obtain neutral phenotypes, we analyzed alignments of endogenous retroviruses (ERVs, see Methods), which are well-annotated ancient pseudogenized copies of retroviruses that are no longer replicating, but are easily identifiable and distributed in large numbers over the human genome[44]. We treat these sequences as independent “replicates” of the evolutionary process and compute the mean and variance of quantitative molecular phenotypes derived from these sequences. We use reconstructed ancestral DNA sequences (based on alignments of closely related primates) along the lineage leading to human, and infer the ancestral phenotypes from these. This allows us to directly study the forward evolution of phenotypes in an evolutionary ensemble without relying on model assumptions for parameter inference.

### Dynamics of GC content and TATA box strength confirm the directional force of mutation

We first considered the “simplest” possible DNA-based phenotype: GC content, a phenotype that has been studied using comparative phylogenetic methods, e.g., [45]. We computed GC content in 100 nucleotide segments extracted from ERVs (see Methods) binned by the GC content of the inferred ancestral sequence. As expected for a simple restoring force of mutation, qualitatively, sequences with relatively high GC content (Figure 1a, triangles) show a decrease over evolutionary time, while sequences that start with a low GC content show an increase in GC content over time (figure 1b, filled squares). As predicted, the decrease at these short evolutionary distances appears linear.

**Figure 1 –.**
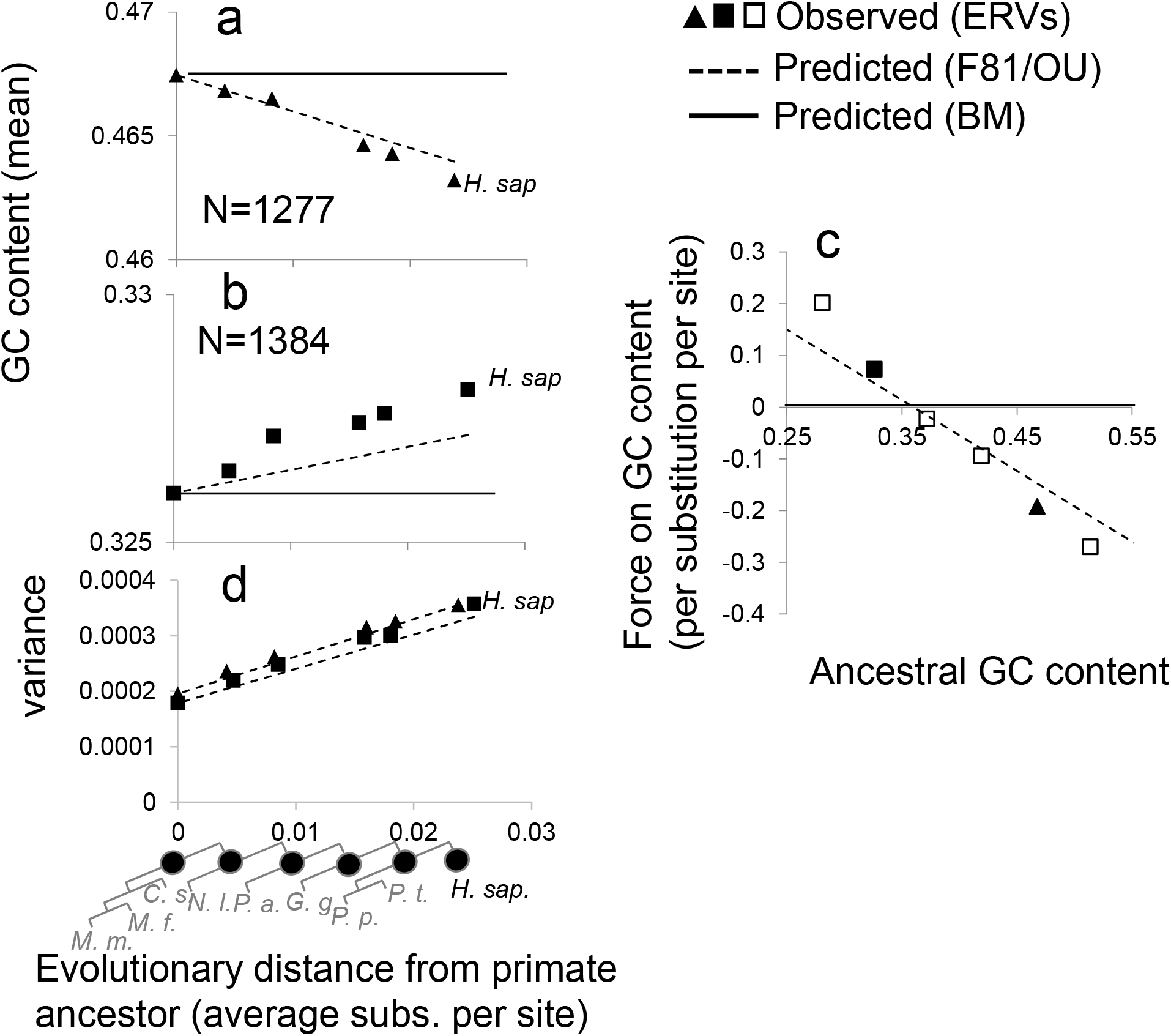
neutral dynamics of GC content. a) observed mean GC content in inferred ancestral primate ERV sequences with GC content in the common ancestor between 0.45 and 0.5 (filled symbols) compared to predictions of the model considered here (F81, dashed line) and the Brownian Motion model (BM, solid line). *H. sap* indicates the extant human sequences. b) as in a), but GC content in the common ancestor between 0.3 and 0.35 (filled symbols) c) inferred strength of evolutionary restoring force estimated by simple linear regression on the GC content for each bin of ancestral GC content (symbols) compared to predictions of the model considered here (F81, dashed line) and the Brownian Motion model (BM, solid line). d) observed variance of GC content in the two bins of ancestral GC content shown in a) and b) (filled triangles and squares, respectively) compared to predictions of an OU model with mean restoring force given by our theory (OU, dashed line). Filled circles below the graphs represent the phylogenetic positions of the reconstructed primate ancestors used in the analysis. *H. sap* indicates the phenotype of the extant human sequences.

To compare quantitatively, we express GC content in the notation of our theory. The GC content of the ancestral sequences in the bin represents *Z*(0) and we used *π* = (*π_A_, π_C_, π_G_, π_T_*) = (0.32,0.18,0.18,0.32) which gives 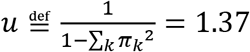. For GC content in a sequence of length L, each locus contributes 1/L if it is G or C, and 0 otherwise. This means 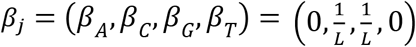 for all *j*. From these we compute the other key parameters for GC content

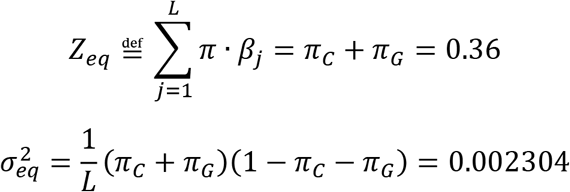

In the Appendix, we show that in the case of GC content, Ψ(X) takes a genotype independent form, i.e., 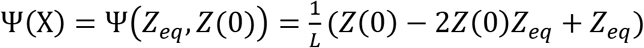, so the dynamics of the variance of GC content can also be predicted from the quantities above.

We found good agreement between the theory based on the F81 DNA substitution model (dashed lines in Figure 1a,b) and the observed changes in GC content. We also used simple linear regression to infer the change in GC content over time as an estimate of the evolutionary force (see Methods). Once again, we found good agreement between the theory and the observations, (Figure 1c) with some slight deviations for sequences with ancestral GC content much different than equilibrium. We believe these deviations are due to the misspecification of our DNA substitution model which does not include CpG bias in the mutation rate or transition-transversion rate bias. Finally, we note good agreement between the variance predicted by the F81 neutral model and the observed increase in variance for GC content (Figure 1d). Given the range of observed GC content and the parameters values above, the rate of increase in variance predicted by the F81 model is within 10% of the prediction of the OU process 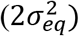, so we lack power to distinguish between these models with these data. This suggests that the OU process (with restoring force set to *u* as defined above) provides a good approximation to the neutral evolution of GC content.

To rule out possible circularity due to the use of probabilistic models of evolution in ancestral genome reconstruction [46], we repeated this analysis using ancestral sequences reconstructed using maximum-parsimony, which makes no assumptions about the relative likelihood of substitutions between different nucleotides. We found qualitatively similar results (Supplementary Figure S1).

Next, we repeated these analyses for the strength of matches to the TATA box ([47], see Methods) in the first 10 residues of each ERV segment. Because positions in transcription factor binding sites contribute approximately linearly to affinity[48], this represents a simple biochemical phenotype whose evolution is well studied[4]. For clarity we note that in our case none of these sequences are expected to have functioning TATA boxes: these are sequences with similarity to the TATA box that arise by chance[49]. The strength of matches to the TATA box is computed using a 4 x 10 weight matrix model, which corresponds to *a* in our model (see Methods). We bin the sequences by the average strength of TATA box in the inferred ancestor, *Z*(0), and again we used *π* = (0.32,0.18,0.18,0.32) which implies *u* = 1.37. The other parameters are *Z_eq_* = −9.223 and 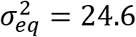, which are obtained directly from the matrix model (see Methods). We found good agreement with the predictions of the model (Figure 2a-c). In this case, we cannot obtain a genotype independent form for Ψ(X), so we do not have a simple prediction for the variance. Nevertheless, once again the linear, constant increase in variance seems to agree reasonably well with the prediction of the OU process (Figure 2d).

**Figure 2 –.**
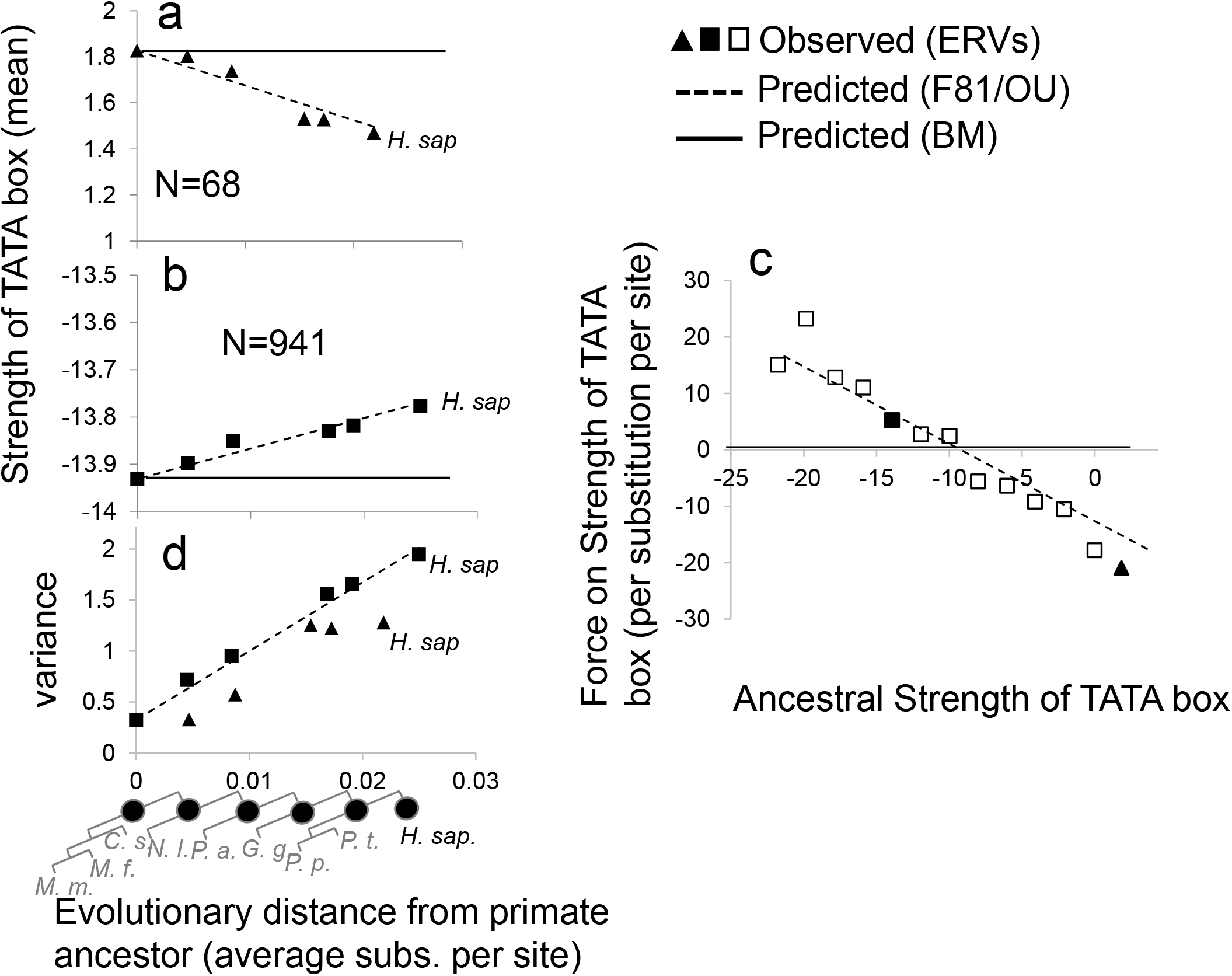
neutral dynamics of TATA box strength. As in figure 1, but for the strength of the match to the TATA box in the first 10 nucleotides of the ERV sequences. Ancestral strength in a) is between 0 and 2, while in b) between −14 and −12.

### Dynamics of ATGs show the predicted mutational force, but no longer match an OU process

We next considered a sequence-based trait where we expected to find example sequences that are far from equilibrium: the number of ATG start codons in 100 bp segments. Sequences can only encode a natural number, 0, 1, 2 … ATGs. Hence even a single point mutation can have a large effect on the phenotype. Once again, we emphasize that we do not expect these ERV sequences to encode functional proteins, so these are simply ATGs that occur by chance or may have functioned in the ancestral viral proteins.

As for GC content and strength of TATA boxes, we found that the number of ATGs in neutral sequences shows a clear mean-reverting force. For example, in ancestral sequences that have exactly 4 ATGs, we see the number of ATGs decreasing over time (Figure 3a, triangles). This is intuitive as we expect random mutation to destroy these “informative” signals over time. Less intuitive is the analogous result for ancestral sequences that start with no ATGs: in these sequences we see (on average) the accumulation of ATGs over evolutionary time (figure 3b filled squares). Evolution appears to be creating start codons, such that overall 142 ATGs are found in human ERV segments that had none in the primate ancestor. Again, we emphasize that the creation of ATGs is not the result of any selection or biological function, rather, in these sequences ATGs are simply being created by random mutations faster than they are being destroyed. We note that the temptation to create adaptive explanations for the appearance of ATGs in these sequences is very strong (see Discussion), but we confirmed that the dynamics of ATGs are similar in simulations where we can be certain there is no selection or function.

**Figure 3 -.**
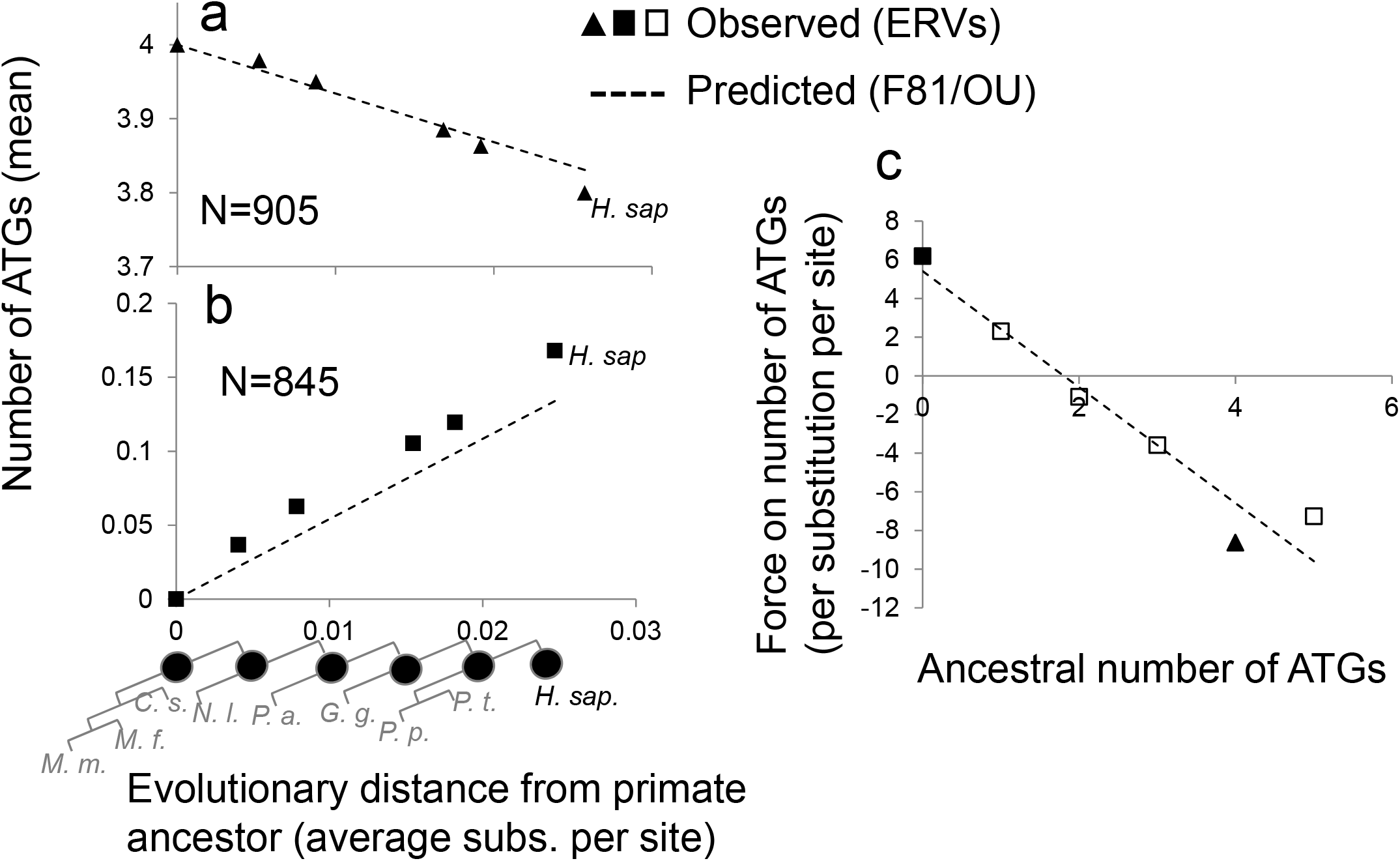
neutral dynamics of mean number of ATGs. As in figure 1, but for the number of ATGs in the ERV sequences. Ancestral number of ATGs in a) is 4, while in b) it is 0.

To predict the dynamics of the number of exact matches to a short sequence, which is not strictly an additive phenotype, we treat the DNA sequences as a series of overlapping w-mers, which we assume are independent; in the case of ATGs, *w* = 3. This means that we imagine 4^*w*^ possible alleles at each locus, and *β_j_* = 1 for the short sequence of interest, and 0 otherwise. If we assume that each DNA letter still evolves independently and at the same per-site rate, the effective evolutionary rate between these alleles will be 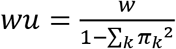. Furthermore, with 4^*w*^ ≫ 1, as long as the mutation process is not too biased, 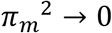, so *u* → 1 and the mutational restoring force is simply *w*. As before, the number of ATGs in the ancestral sequences in the bin represents *Z*(0). The equilibrium phenotype in this model can be computed exactly

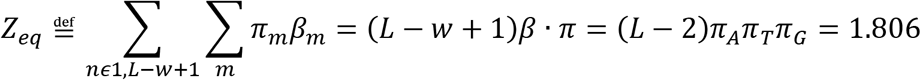

As with GC content and strength of TATA boxes, we find very good agreement between the theory and the mean dynamics (dashed lines in figures 4a,b,c) and clear evidence for the linear restoring force with strength simply equal to 3 start codons per substitution per 100nt sites (Figure 3c). We note that the agreement between the observations and theory must be approximate in this case, because the exact dynamics of the number of ATGs is strongly genotype dependent: initial sequences with many “one-off” sequences (e.g., ATC, ATT, ATA, AAG, etc.) are much more likely to increase their number of ATGs than sequences with few of these “one-off” sequences. We believe that because we are measuring the average over a large number of initial sequences, these effects are averaged out (see Discussion).

**Figure 4 –.**
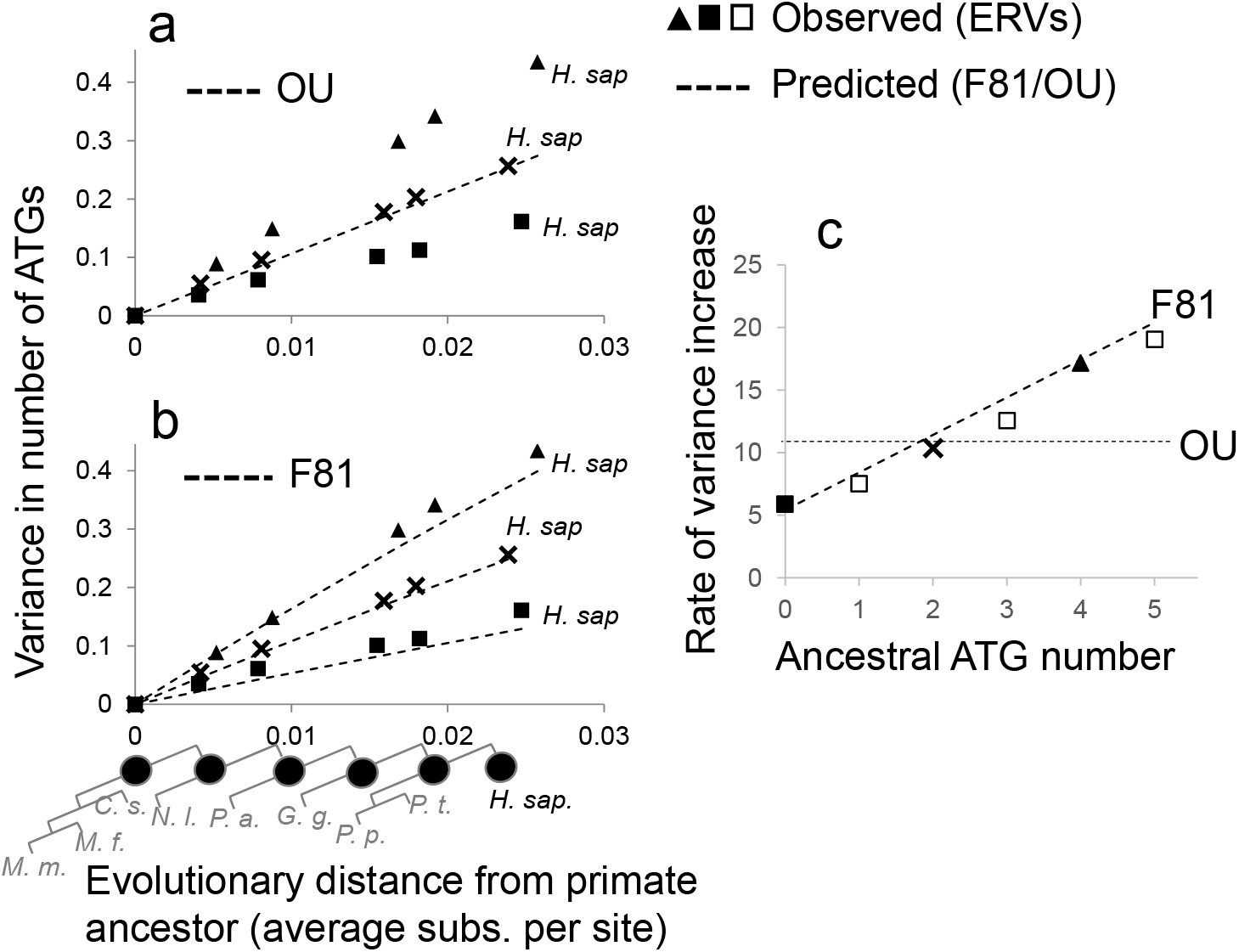
neutral dynamics of the variance of ATGs. a-b) variance of number of ATGs in 100 bp ERV segments that in the primate ancestor are inferred to have no ATGs (filled squares), 2 ATGs (x’s) or 4 ATGs (filled triangles). Predictions based on an OU process model and the F81 model developed here are shown as dashed lines in panels a) and b), respectively. c) the rate of increase of the variance (inferred using simple linear regression) as a function of the ancestral ATG number (symbols) compared to the predictions of the OU model or the F81 model developed here.

Since the number of exact matches is approximately binomial, the long-time variance will be approximately

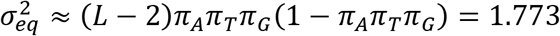

Unlike for GC content and strength of TATA boxes, we observed that the variance in number of ATGs was not independent of the starting phenotype *Z*(0), and therefore inconsistent with the OU prediction (dashed line in figure 4a). We found that sequences with more ATGs than the equilibrium (4 ancestral ATGs, triangles in figure 4a) showed a faster increase in variance than sequences with fewer (0 ancestral ATGs, filled squares in figure 4a). We therefore looked at the ancestral bin that was closest to the mutational equilibrium (2 ancestral ATGs, x’s in figure 4a) and found that the variance increased in very good agreement with the OU prediction, consistent with our theoretical prediction that the F81 model matches the OU dynamics when the phenotype is at equilibrium (Appendix).

Our approximate model for the number of ATGs described above corresponds to a simple phenotype, which is just the counts of the k-th allele [30]. For this phenotype, like for GC content, the F81 neutral dynamics of the variance are independent of the initial genotype, X, but they are still more complicated that the OU model predicts (see Appendix). In this case, Ψ(X) ≈ *Z_eq_* + *Z*(0), with the approximation valid for *π_k_* ≪ 1. At short evolutionary distance, and since for ATGs, *π_k_* ≪ 1, we obtain a simple approximation for the variance of number of ATGs under the F81 model:

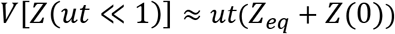

To our knowledge this simple prediction for the rate of increase of variance based on the initial phenotype has not been reported under the house-of-cards model[22]. Comparing this to our observations for the variance of ATGs (with *u*=3) gives remarkably good agreement, and better predicts the linear increase in variance than the OU model (Figure 4b,c). Taken together, these results confirm the prediction above that the true dynamics for molecular phenotypes do not match an OU process when the phenotype is strongly sufficiently far from the mutational equilibrium.

### Neutral dynamics of more complex molecular phenotypes also show a linear restoring force

Many of the molecular phenotypes of interest (e.g., gene expression level) are much more complicated than GC content or strength of binding and are hard to express as additive phenotypes. Nevertheless, a recent study of the mutational effects on gene expression used simulations parameterized by empirical measurements to show simple linear dynamics of gene expression phenotypes[50] even though the traits are unlikely to be linear. To test the generality of our finding of a mutational force on the mean phenotype, we next empirically studied the neutral dynamics of non-additive phenotypes that can be computed from sequence. We chose three phenotypes that are of more biological interest: the number of TATA boxes in a 100 bp sequence, the length of the longest encoded peptide, and the intrinsic disorder in the longest encoded peptide. The first of these relates to how non-coding sequences evolve so-called homotypic clusters of binding sites[51], [52], and the latter two of these phenotypes are related to the emergence of new protein-encoding genes from random DNA [53], [54]. To obtain these, we computed the lengths of six-frame translations from our 100-basepair ERV segments, and the propensity of the peptide to fold ([55], see methods). We emphasize that these phenotypes are highly non-linear in the DNA sequence genotype, and therefore strongly violate the assumptions used to derive our results above.

Remarkably, we found that these phenotypes also showed simple linear mean dynamics, albeit with stronger evolutionary forces pushing the phenotypes to their mutational equilibrium (Figure 5a) than the additive phenotypes considered above. The force appears to be different for each phenotype, but we have no theory to predict it. To a large extent, the evolutionary force appears simply proportional to the distance of the ancestral sequence from the equilibrium (Figure 5b) and the quantitative strength of the force is less than 10 units per 100 bp DNA sequence. Thus, non-additive phenotypes also appear to show a simple force of mutation proportional to the distance of the ancestral phenotype from the mutational equilibrium.

**Figure 5 –.**
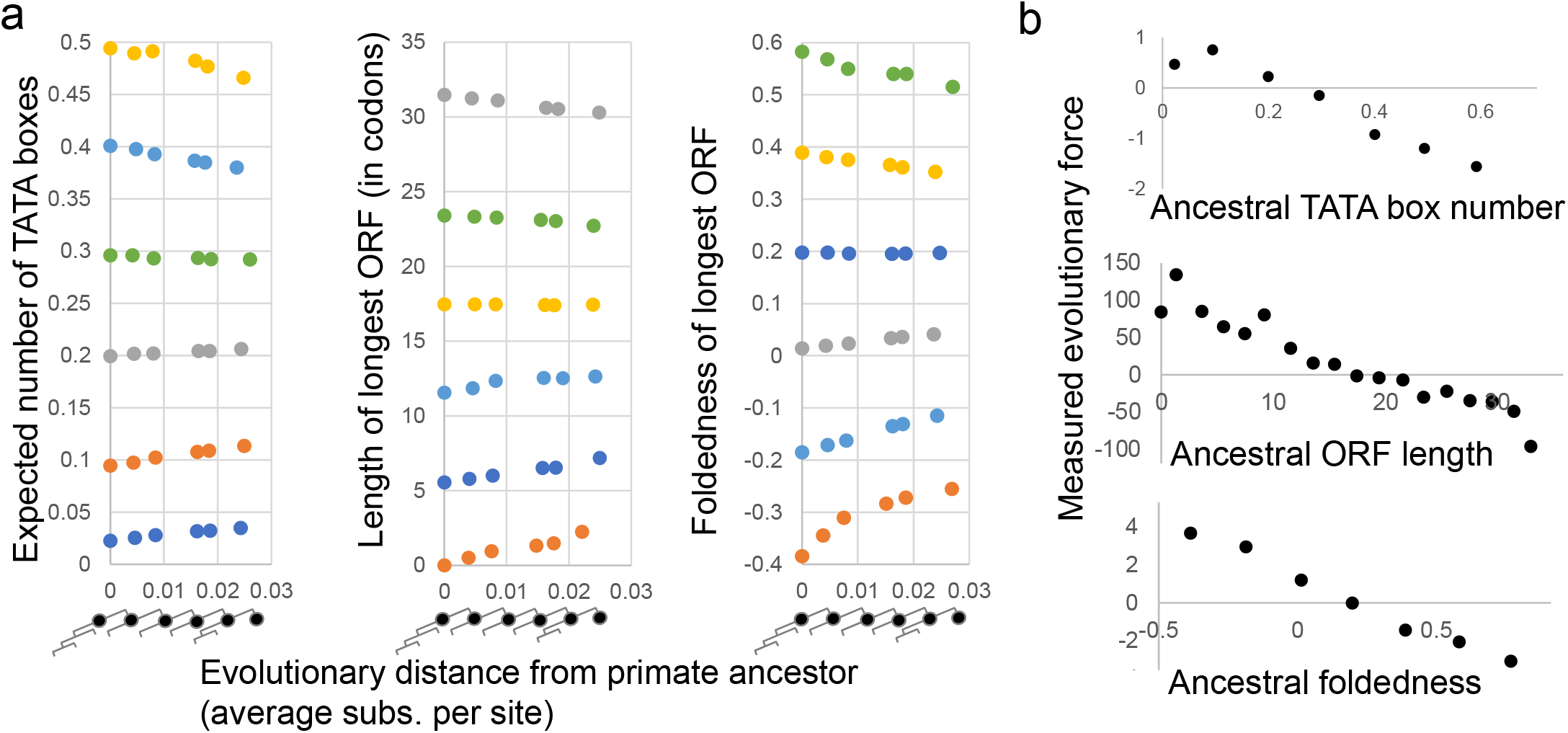
mean dynamics of non-additive molecular phenotypes show mutational restoring force. a) three molecular phenotypes computed from 100 bp ERV segments in reconstructed sequences, binned by their values in the primate ancestor. Colours indicate different ancestral values. See text for phenotype details. b) inferred mutational restoring force as a function of ancestral phenotype, estimated through simple linear regression.

### The BM null hypothesis cannot always be rejected using standard inference approaches on GC content in ancient ERVs

To test whether the predicted and observed deviations from the Brownian Motion (BM) null hypothesis were strong enough to be detected in a typical comparative phylogenetic analysis, we implemented standard Gaussian maximum likelihood inference ([13], described in the Appendix; experiments using the OUCH package [43] for inference did not converge reliably, and led to errors in estimation). We obtained six well-aligned fragments from an alignment of 39 mammals (see Methods, supplementary Figure S2) for an unusually ancient ERV that is thought to evolve in the absence of selection[56]. We found that corrected AIC was the lower for BM than for OU for all 6 segments (Table 1) because the likelihood of the data under BM and OU was very similar, so the simpler model was chosen. We also implemented maximum-likelihood inference for the F81 neutral model (See Appendix and Methods), and compared it (with parameters set to match the GC content of the ancient ERVs) to BM. We found that it was favoured for two of the 6 segments, BM favored in another two, and the other two showing similar corrected AICs. These results suggest that despite theoretical and empirical evidence for a mutational force, in an inference set up typical of phylogenetic comparative analysis, the BM model, which does not include this force, fits the data as well as models (OU and F81) that do include this force. We note that these results cannot be explained by noise [42]in our phenotypic measurements, as GC content is computed exactly from sequences.

**Table 1 –.** differences in corrected AUC between OU, F81 and BM. Each cell represents the corrected AIC comparison vs. the Brownian Motion (BM) null hypothesis. AIC is inversely proportional to likelihood, so lower corrected AICs represent better fitting models.

| Segment | Observed ERV segments |  | Simulated ERV segments |  |
| --- | --- | --- | --- | --- |
|  | OU | F81 | OU | F81 |
| 10270_10563 | 2.35 | -2.27 | 2.15 | -0.80 |
| 11935_12275 | 2.46 | 0.63 | -29.0 | -3.94 |
| 11196_11705 | 2.39 | -1.94 | -6.57 | -4.74 |
| 320_722 | 2.36 | 2.57 | 2.40 | -9.90 |
| 7916_8695 | 2.36 | 0.57 | -13.8 | -3.38 |
| 8938_9248 | 2.37 | 2.60 | -18.5 | -6.67 |

Although this ERV is thought to evolve in the absence of selection[56], our prediction of the dynamics of the mean and variance of GC content is based on a simple DNA substitution model, which only captures some features of sequence evolution. We wondered if mis-specification of the neutral model could explain the lack of evidence for mutational force in the ancient ERVs. We therefore simulated molecular evolution of these ERV sequences under a simple neutral model of DNA substitutions (see Methods) and found much clearer support for models that do include the force of mutation: for the simulated data, 4/6 segments favour OU over BM, and 5/6 favour the F81 neutral model over BM (Table 1). These results confirm that simple substitution models do produce a measureable force of mutation, and suggest that there are other aspects of the neutral evolutionary processes in the real ERV sequences that is not accounted for by either OU or F81 models, but is better captured by Brownian Motion (see Discussion).

## Discussion

Our results show that mutation is expected to aid the creation of molecular phenotypes *de novo*[28] and relate to several areas of research in molecular evolution. For example, in agreement with simulation results [57], we find that mutation alone is sufficient to create transcription factor binding sites in DNA sequences, and we quantify this effect: 100 nt sequences that have no strong matches to the TATA box are expected to accumulate them at a rate of approximately 0.5 TATA box matches per substitution per site. Similarly, in random 100 nt sequences with no open reading frames, mutation will tend to create open reading frames at a rate of >50 codons per substitution per site. Even if the encoded peptides start out as strongly disordered[54], the force of mutation is expected on average to increase their tendency for folding[53]. Remarkably, but consistent with a recent report [35], even though all of these molecular phenotypes are non-additive, they show simple linear change over time, following our predictions for additive phenotypes. We believe that this is because our ERV sequences are relatively near the mutational equilibrium for these phenotypes, and speculate that the evolutionary dynamics are approximately linear near the mutational equilibrium. It will be of great interest to extend the theoretical work to explain these observations.

Unlike previous theoretical work which focused on quantifying the force of mutation on phenotype evolution by including mutational bias[33], [58], here we used the F81 substitution model, which is formulated in terms of the (possibly non-uniform) equilibrium frequencies for the DNA bases (alleles). Our results suggest that the mutational equilibrium phenotype, rather than the mutation rate bias, plays the key role in determining the neutral dynamics of phenotypes, consistent with recent theoretical results[30], [33]. Although the F81 model does have mutation bias (the rate bias from allele m to allele k is π_m_/ π_k_; there is no transition-transversion bias or CpG hypermutation), we also considered the Jukes-Cantor model[20], which is a specific instance of the F81 model where all π_m_=1/4 (u=4/3) for DNA, and therefore shows no mutation bias. Importantly, even under this simplification, we still predict a restoring force proportional to the distance from the mutational equilibrium, although some simplification of the dynamics is obtained (e.g., the neutral variance dynamics of GC content is no longer genotype dependent). Even simplified bi-allelic (non-DNA based) models with no mutation bias show a restoring force of mutation when far from mutational equilibrium (Appendix), strongly suggesting that the force of mutation in phenotype evolution is not due to mutation rate bias, but is a more fundamental result of the mapping between the discrete genotype space and the continuous (or ordinal) phenotype. We did not pursue these simplified models further because the predictions of the dynamics were not close to our observations from ERV sequences, presumably because the predicted mutational equilibrium is too far from the real data (e.g., equilibrium GC content of 50%, rather than 36%). This highlights the need to develop simpler (approximate) theory that can still be quantitatively compared to data.

The dynamics of the mean phenotype in the F81 model are similar to those expected under the house-of-cards mutation model[22], a model defined and derived using only phenotypic quantities. The similarity is most obvious if the mean of the phenotype effect distribution in the house-of-cards model is taken to be the “mutational equilibrium” phenotype, *Z_eq_*, defined above. To understand how this similarity arises, we note that if the F81 DNA substitution model is viewed as a (4-dimensional) phenotype model, the dynamics of the genotype probabilities match the mean dynamics of a house-of-cards mutation model. Since the mean of an additive phenotype is a linear combination of the genotype probabilities, it follows the same dynamics.

For GC content and strength of matches to TATA boxes we found agreement with the predictions of the variance dynamics based on the OU process, even though we can show that this is only approximate (See Appendix). We believe this is because we measured these phenotypes in ERVs that are close enough to mutational equilibrium, where we expect the dynamics to be well approximated by the OU process. However, neutral phenotype evolution models (such as the F81 model developed here) should accommodate phenotypes that may have been under strong selection pressure in the past, making it unlikely that the initial ancestral phenotype is close to mutational equilibrium at the time selection is relaxed. This scenario is consistent with observations from mutation accumulation studies[26], where once selection is weakened, phenotypes evolve in a directional manner.

Despite the clear empirical evidence for the force of mutation on GC content at short evolutionary distances when we averaged over thousands of sequences, we found inconclusive results when comparing models using a standard maximum likelihood/AIC inference analysis of GC content in ancient ERVs. In our experiments Brownian Motion, which does not include the force of mutation, fit the data as well as models that include the force of mutation. We speculate that a considerable portion of the variance in GC content at longer evolutionary distances is due to neutral processes not included in simple DNA substitution models. Understanding these sources of variance is an area for further research.

More generally, the similarity of the dynamics of neutral phenotypes to an OU process suggests that at time-scales considered here, similarity of phenotype evolution to OU process should not be used as evidence for purifying selection (as suggested in[1]). Further, since even neutral phenotypes show more complicated variance dynamics (genotype-dependent and nonmonotonic) than the OU model predicts, our results add the possibility of misspecification of the underlying models[59] to the previously reported challenges with phylogenetic inference of OU models[42], [43], [60]. Developing tests based only on the mean phenotype (which does seem to match the OU process assumptions remarkably well) is an area of promise, though it remains unclear how much information about mean dynamics is contained in the observations at the tips of the tree[60]. On the positive side, our results are consistent with the idea that mutation and selection act in a simple additive way on the evolution of mean phenotypes[11], [30], both leading to OU-like dynamics. Further, our results limit the quantitative range of restoring force expected due to mutation alone: if the restoring force estimated in an OU model exceeds the predicted value, we believe this can be interpreted as stabilizing selection. Similarly, large changes in the optima in multi-optimum OU models [10] are unlikely to result from mutation alone.

Finally, our approach of using reconstructed ancestral sequences to empirically study the forward evolution of phenotypes is applicable whenever the genetic architecture of a trait is known. With the increasing power of association and other high-throughput studies to determine the loci and their effect sizes for many phenotypes[61], [62], additive models of traits can be inferred. Furthermore, machine learning methods trained on genome-scale data and massively parallel assays may be able to predict non-additive molecular phenotypes from sequences [63], [64]. Once a genotype to phenotype model is defined, observed phenotype evolution can be compared directly to neutral phenotypes obtained from neutral sites (such as ERVs[44]) or simulations at the sequence level[65]. The simulation approach has been applied successfully to molecular traits that can be computed directly from intrinsically disordered protein sequences[66], [67]. If applied more generally to quantitative characters, this could be viewed as an extension to practice of inferring phylogenetic trees from genotypes, even when studying phenotypes [18], [24]: the null hypotheses for phenotype dynamics can also be obtained using our understanding of genotype evolution, further cementing the inevitable merger of systematics and evolutionary genetics[18].

## Methods

### Unconstrained sequences in primate genomes

We obtained 6362 coordinates for predicted ERVs in the human genome from the gEVE database[68], corresponding to a subset of ERV segments that span more than 100 nucleotide residues and contain no ambiguous letters. We obtained alignments of these along with their reconstructed ancestors[46] from the 100 vertebrate alignment tree from Ensembl, we extracted the species in the aligned segments using the Ensembl compara REST API[69]. As a control for possible circularity due to probabilistic models of sequence evolution used in ancestral genome reconstruction [46], we also performed analysis on ancestral sequences reconstructed using the maximum parsimony criterion as implemented in the phangorn package[70].

We used the extant human sequence, as well as the following reconstructed ancestors using the naming convention from ensembl: Hsap-Ppan-Ptro[3], Ggor-Hsap-Ppan-Ptro[4], Ggor-Hsap-Pabe-Ppan-Ptro[5], Ggor-Hsap-Nleu-Pabe-Ppan-Ptro[6], and Csab-Ggor-Hsap-Mfas-Mmul-Nleu-Pabe-Panu-Ppan-Ptro-Tgel[11] (square brackets in these names refer to the ancestor number in the ensembl tree are not references). For each ERV alignment we extracted segments of at length 100 with <10% gapped columns, required all sequences to be present in each alignment, yielding 8386 segments. Csab-Ggor-Hsap-Mfas-Mmul-Nleu-Pabe-Panu-Ppan-Ptro-Tgel[11] was considered the “primate ancestor” and the value of the phenotypes computed based on this sequence was used as the ancestral phenotype, Z(0). Binning was done based on this value, and each of the descendent sequence phenotype values were assigned to this bin. Phenotype mean and variance were calculated for all sequences in the bin, and these sample sizes are reported as N for bins shown in figures 1, 2 and 4.

For each “descendent” sequence, we computed the evolutionary distance in substitutions per site from the ancestor using the Juke-Cantor model and used the average of these as the evolutionary distance for that phenotype bin. The common ancestor was always assigned a distance of zero.

To estimate the mutational restoring force and rate of variance increase, we used simple linear regression to estimate the slope of the regression of the mean phenotype and variance of the phenotype on time (in substitutions per site). Because the ancestral value was determined by the binning process, and the evolutionary time for the ancestor was zero by construction, to avoid potentially biasing the estimate of the slope, we excluded the common ancestor from the regression and used the remaining 5 datapoints. R^2^ for these was typically >0.9.

### Strength and number of TATA boxes

We obtained the matrix model of TBP from the Jaspar database [47].

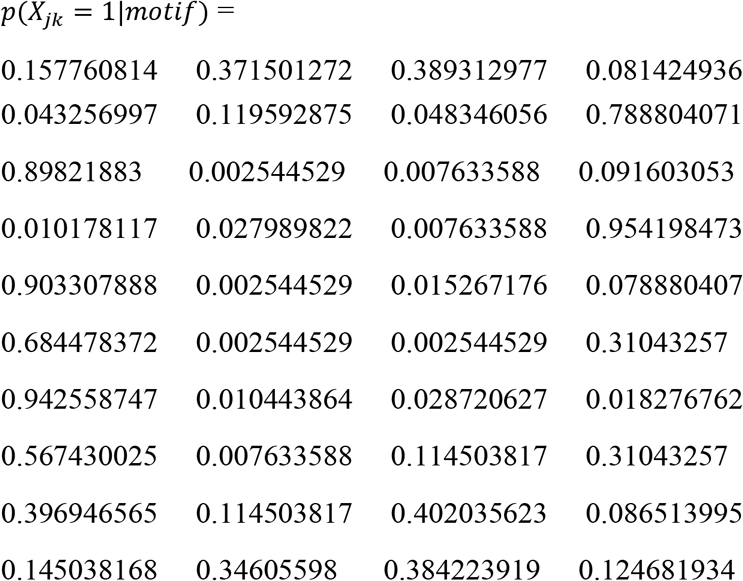

Where *p*(*X_jk_* = 1|*motif*) is the probability of observing the kth allele at the jth position in the transcription factor binding probability matrix.We assumed a position independent background model, as above, *π* = (0.32,0.18,0.18,0.32), so we define 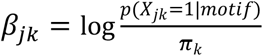. For the strength of TATA box analysis, we used the strength of the match in the first 10 bp of each ERV alignment and used the standard log-likelihood ratio score, S, as a measure of binding strength[71]. To compute the other parameters for the strength of TATA boxes, which have width w=10, we used

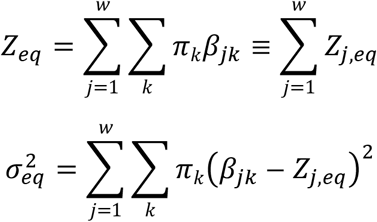

For the expected number of TATA boxes in the entire sequence, we used the sum of the posterior probability of each position being a match to the motif[71], 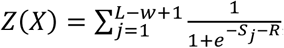, where each *S_j_* represents the standard likelihood ratio score [71] for the subsequence of length w starting at position j, and *R* is the (position independent) log odds of the prior probabilities of observing a match to the motif or not at each position. 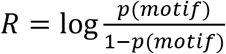. We used 1/100 for this prior, so 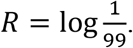.

### Alignment, phylogeny, simulation and inference for ancient mammalian ERVs

we retrieved the human ERV sequence [56] and used BLAST on the Ensembl database[72] to obtain an alignment with all orthologous mammalian sequences therein. After filtering the results for quality, we were left with orthologous matches in a total of 39 other mammalian species. We manually extracted 6 segments with relatively few gaps and most of the species present (supplementary data).

For each of the 6 segments, we used PAML (v4.9) [73]to infer the branch lengths of the evolutionary tree and reconstruct the corresponding ancestral sequence of the 40 mammalian species included in the study. Then, we employed a DNA substitution simulator [65] to evolve the reconstructed sequences along the mammalian phylogeny. For each set of real or simulated sequences we computed the quantitative molecular phenotypes. Tree was drawn using plot function from the APE package for R [74].

We implemented maximum likelihood inference (described in detail in the Appendix) for the three models of phenotype evolution considered here (BM, OU, F81) in python using scipy [75]. To numerically optimize the log-likelihoods (log *L*), we used Broyden–Fletcher–Goldfarb–Shanno (L-BFGS-B) starting with random initializations of parameters. We confirmed the inference results comparing the likelihoods of BM and OU models to those obtained with the Geiger package[76]. Models were compared using corrected AIC (AICc), which is defined as 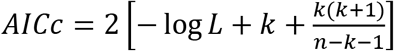, where *k* is the number of parameters and *n* is the number of species for which we have a mean phenotype measurement.. We adopted the widely-used assumption for inference using OU models in comparative phylogenetic analysis that the OU process is initially at equilibrium[76]. Models with corrected AIC ~2 units below the competing model was deemed better fitting.

Alignments, scripts and evolutionary trees are available at https://github.com/shz9/neutral-phenotype-model.

### Foldedness of longest peptides

We considered all 6-frame translations within the 100bp sequences starting with ATG, and selected the longest. Using these peptides, we measured the propensity to fold using the method described in [55]. In this scale, positive values indicate folded, while disordered regions obtain negative values.

## Acknowledgements

We acknowledge Dr. Amin Zia, Gavin Douglas, Caressa Tsai, and other members of the Moses Lab who contributed ideas and valiant attempts to this research over many years. We acknowledge Prof. Michael Laessig for hosting SZ during part of this research. We thank Prof. Michael Lynch for helpful comments on the manuscript. AMM and SZ were supported by NSERC discovery and CFI grants to AMM.

## Supplementary figures

**Figure S1 –.**
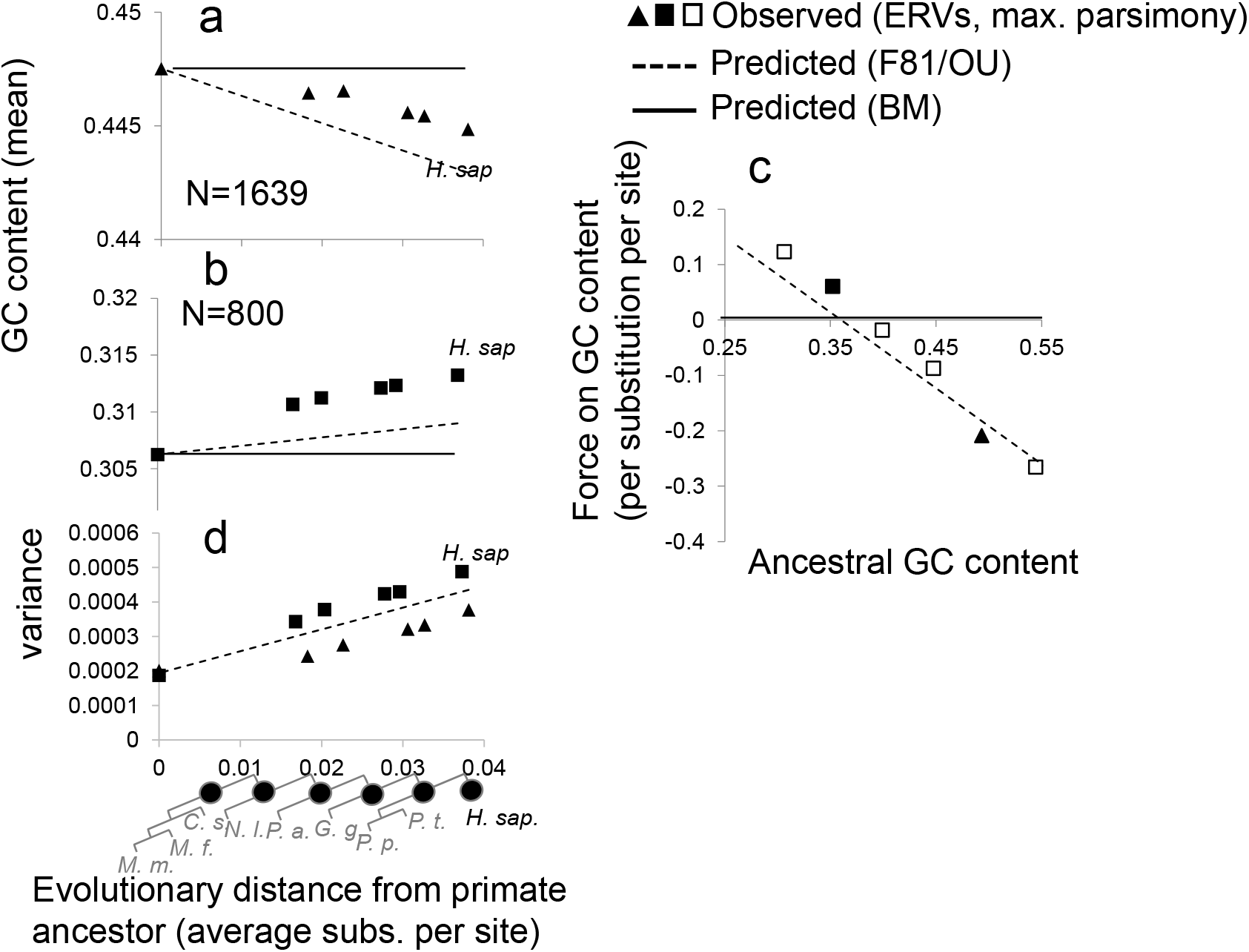
as in figure 1a-c, but with ancestral reconstruction done using maximum parsimony

**Figure S2 –.**
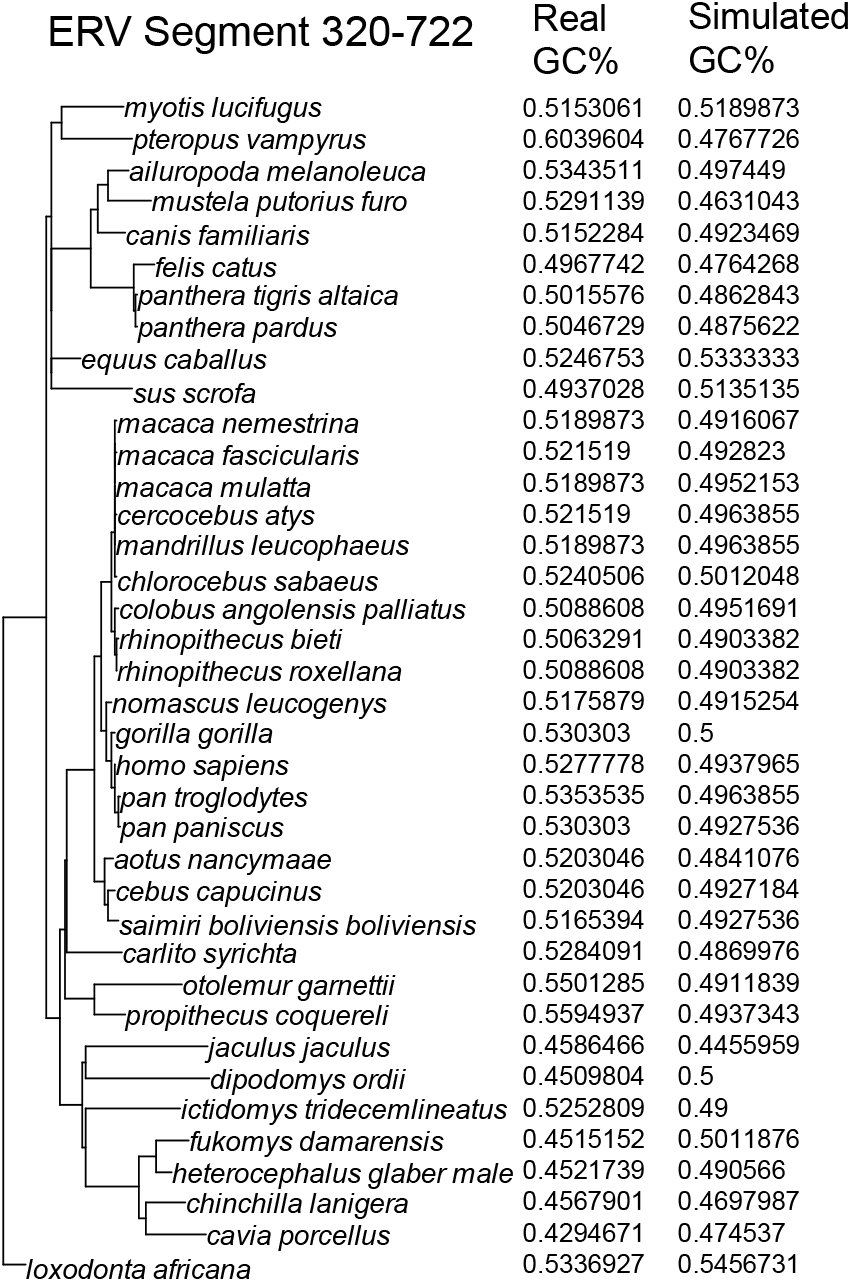
example phylogeny and GC content in ancient ERVs. The inferred phylogenetic tree and associated GC content measurements (GC%) for the segment 320 – 722 for the species used in the Maximum Likelihood inference analysis of GC content.

**Figure S3 –.**
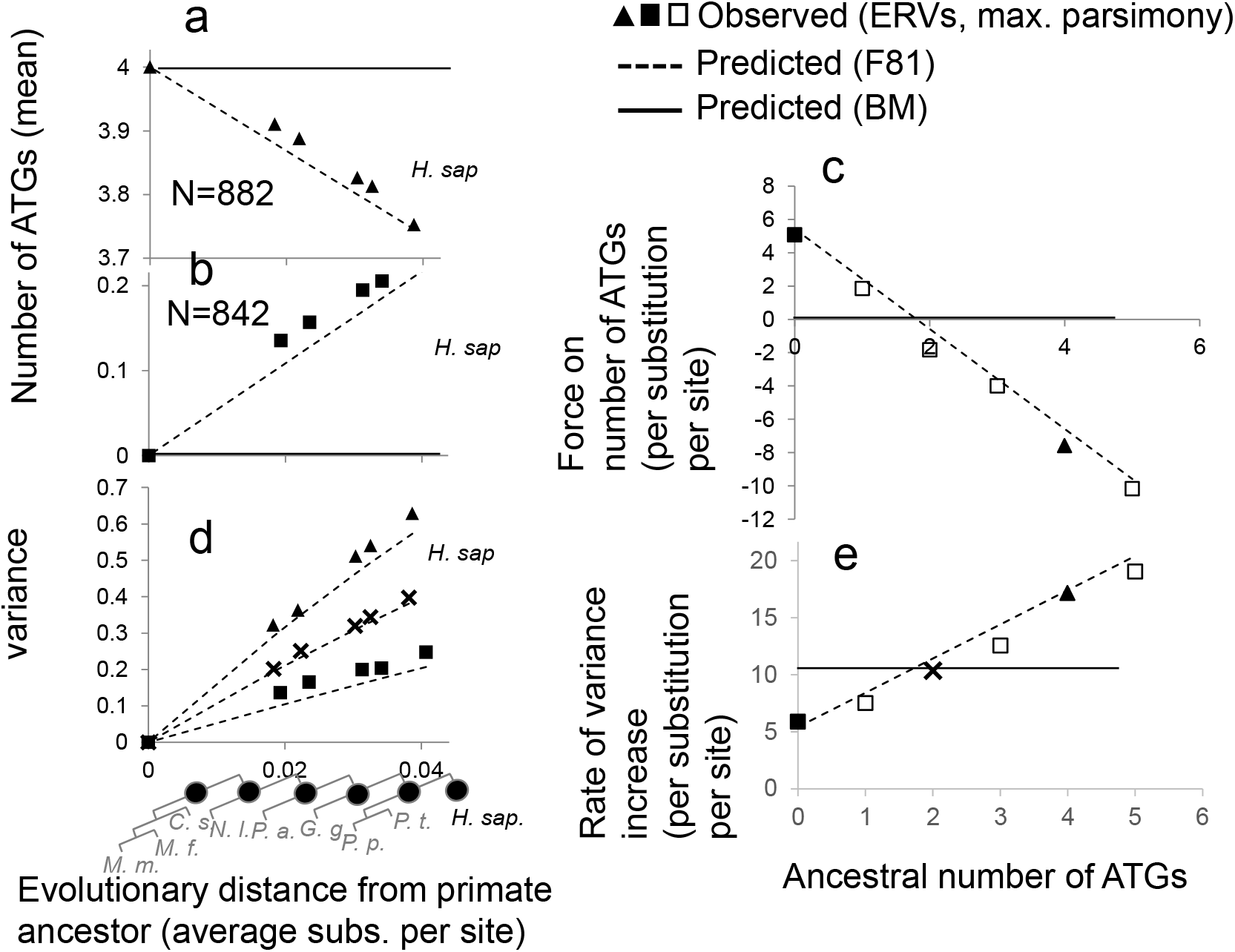
as in figure 3 and 4, but with ancestral reconstruction done using maximum parsimony

## Appendix

## A bi-allelic model

To understand the evolutionary dynamics of mean phenotypes, we start with a linear additive trait that is expressed in a haploid organism with a bi-allelic genetic architecture. Assuming that this trait is underpinned by *L* bi-allelic loci, using the notation commonly employed in modern GWAS literature, we can express the dependence of the phenotype on the underlying genetics as follows:

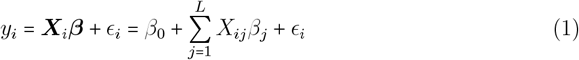

Here, *y_i_* denotes the phenotype for individual *i*, ***X***_*i*_***β*** is the genetic contribution to the phenotype and *ϵ_i_* captures the contribution of the environment and other non-linear effects. ***X***_*i*_ is a 1 × (*L* + 1) binary vector indicating whether individual *i* has the reference or alternative alleles at each locus *j*:

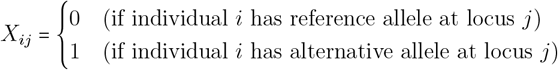

***β*** is an (*L* + 1) × 1 vector of effect sizes. We assume, without loss of generality, that at each locus, the effect sizes for the reference and alternative alleles are encoded by *β*_*j*1_ and *β*_*j*2_, respectively. This implies that *β_j_* = *β*_*j*2_ – *β*_*j*1_ and from this, it follows that the intercept term *β*_0_ is the sum of the effect sizes of the reference alleles at each locus: 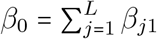 (e.g. individuals with no mutations at any site will have a genotypic value of *β*_0_). Modern GWAS methodologies typically dispense with the intercept term by standardizing the phenotype vector and genotype matrix. In our case, it is a constant that will not affect the dynamics of phenotypic evolution and in fact has a value of zero for many of the traits that we will consider in the main text.

The main quantity of interest in our analysis is **the mean phenotype in a population or species** 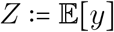. If we assume that the effect sizes are fixed, we obtain:

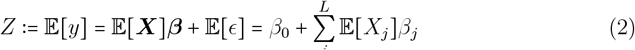

The environmental contribution is omitted because we follow the standard assumption that 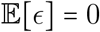. Furthermore, in the context of this analysis, most of the traits that we are interested in are expressed at the molecular and cellular level, often comprising aggregate statistics of the underlying sequence, e.g. GC content. Therefore, the environmental contribution is assumed to be nil. Note that within a given population or species, 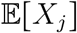 is simply the allele frequency *f_j_* at the *j^th^* locus.

In Phylogenetic Comparative Methods [7, 6], one of the central aims is to analyze the evolution of the mean phenotype over time *Z*(*t*), given an initial starting point *Z*_0_:= *Z*(0). In particular, given an initial mean phenotype, the goal is to characterize its evolution over an ensemble of different hypothetical populations or experimental replicates. Assuming that the effect sizes (*β_j_*s) and the genetic architecture of the trait remain constant, the only time-dependent parameter is the genotype at each locus:

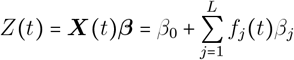

**Figure 1:**
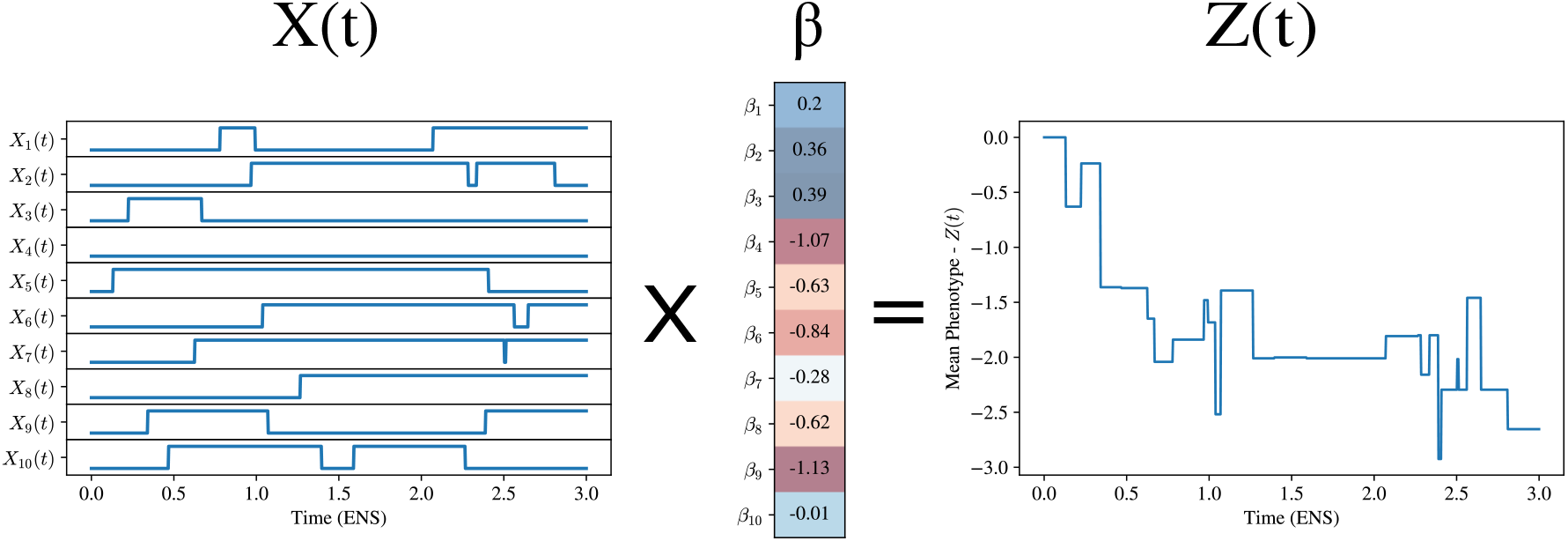
An illustration of the bi-allelic model for the evolution of mean phenotype in a single population. In this example, we show the allelic state of *L* = 10 genetic loci as a function of time ***X***(*t*), measured in units of Expected Number of Substitutions (ENS), and the corresponding effect sizes ***β***. In this simulation, all loci start with the reference allele and then, through mutation and drift, alleles can undergo fixation or extinction. We model this evolutionary process with a substitution model where loci evolve according to a Continuous-Time Markov Chain (CTMC). The mean phenotype as a function of time *Z*(*t*) is obtained by multiplying the state of the locus at time *t* with the effect sizes according to Equation 3.

Characterizing the evolution of the genotype matrix ***X***(*t*) over time in response to mutation, selection, and drift has been a central focus of population genetics theory for the past century [6]. Much of the recent effort in this area made use of the diffusion approximation, first introduced in the works of Wright and Fisher and then later extended by Kimura, among others [4]. In this work, we are interested in characterizing the evolution of the genotype matrix in response to mutation and drift alone and over phylogenetic timescales. In this context, we make use of 2 simplifying assumptions: (1) The loci evolve independently and (2) Evolution proceeds in the weak mutation regime, where at any point in time the population is largely monomorphic. In practical terms, this means that instead of modeling the full trajectory of *f_j_*(*t*) from introduction to fixation or extinction, we model the state of the locus as a jump process between 2 states (reference vs. alternative allele), governed by a Continuous Time Markov Chain (CTMC) [14].

At the risk of overloading notation, for simplicity we replace the continuous random variables 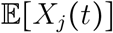 and *f_j_*(*t*) with a binary random variable *X_j_*(*t*) that captures the state of the locus *j* in a given population at any point in time *t*:

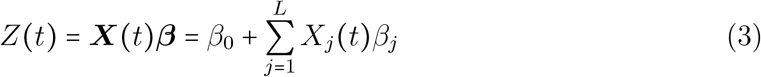

This model is illustrated in Figure 1. Since the phenotype model is linear with respect to the individual loci, most of the quantities that we need to study the evolutionary dynamics of mean phenotypes can be derived as simple functions of single-locus statistics, which we turn to in the next section.

### 1.1 Single locus statistics

As explained in the previous section, we model the genotype *X_j_*(*t*) at a given locus *j* with a Continuous Time Markov Chain (CTMC) that jumps between 2 states corresponding to the 2 different alleles. Because of the way we setup the phenotypic model, the states of the system in this case are *X_j_*(*t*) = 0 if the reference allele is fixed at time *t* and *X_j_*(*t*) = 1 if the alternative allele is fixed. Under this model, the reference allele is expected to be substituted by the alternative allele with an infinitesimal rate λ and the converse happens with rate *κ*λ, where *κ* > 0 is a parameter that quantifies the extent of mutational bias. The generator matrix *Q* of this substitution process has the following form:

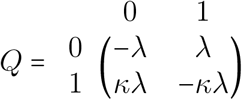

From the above generator matrix, it is straightforward to obtain expressions for the transition probabilities in a time interval *t*:

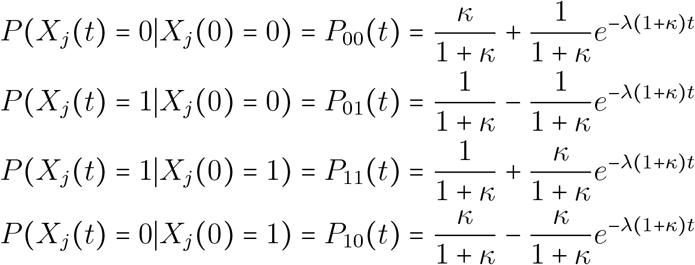

The transition probabilities defined above provide the probability of transitioning from one state to the other in a given time interval. To obtain an expression for the probability that the alternative allele is fixed at any point in time, we marginalize over the ancestral state:

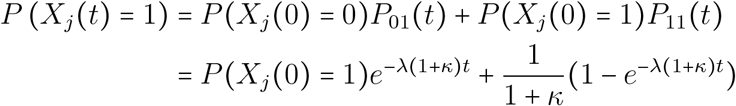

The complement of this, the probability that the reference allele is fixed at any point in time is *P*(*X_j_*(*t*) = 0) = 1 – *P*(*X_j_*(*t*) = 1). If we take the limit as *t* → ∞, we see that this system has a stationary distribution with 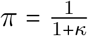 and 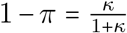. We therefore write the probability of observing the alternative allele at time *t* as:

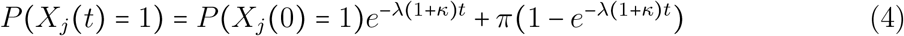

Before proceeding, it is worth highlighting that in this context we use probability in the frequentist sense: we envisage a large ensemble of populations emanating from a single founder population, as in a phylogeny or a set of genes that evolve independently after a duplication event [6]. Each population may lose or fix alleles independently and we wish to characterize the dynamics of mean phenotype over this hypothetical ensemble (see Figure 2). This is in line with the tradition in the Evolutionary Quantitative Genetics literature where researchers model the phylogenetic (i.e. cross-population) mean and variance of the mean phenotype.

Next, in order to derive expressions for the evolutionary dynamics of mean phenotypes, we need to derive expressions for the moments of allelic state at each locus. These quantities will be used in later sections to derive the phylogenetic (i.e. ensemble) mean and phylogenetic covariance for the mean phenotype.

**Figure 2:**
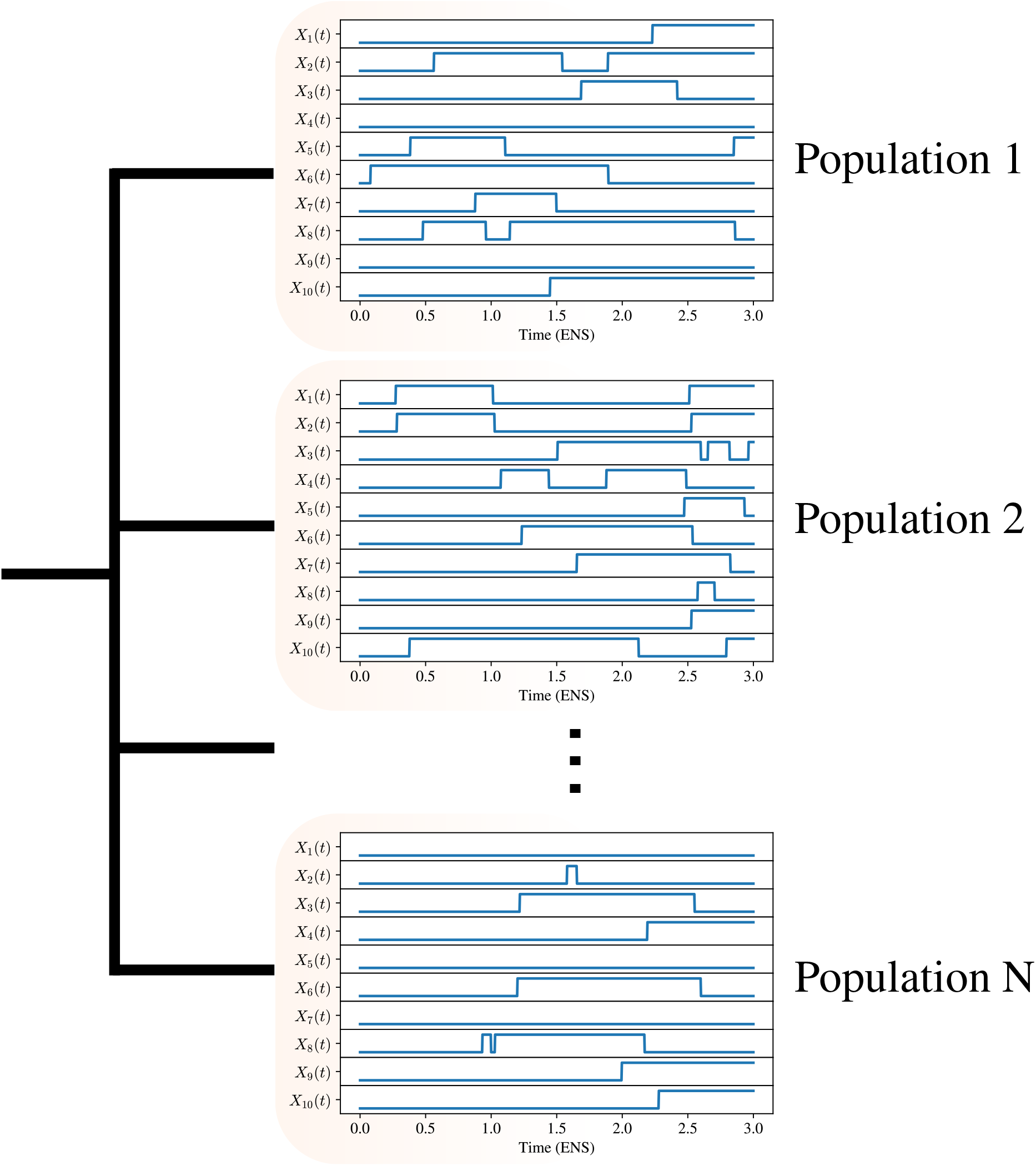
An illustration of the evolutionary process in an ensemble of *independent* populations or experimental replicates. At time *t* = 0, a single founder population gives rise to an ensemble of *N* separate populations that are then presumed to evolve completely independently. In each population, the (bi-allelic) loci are subject to mutation and drift, which can result in the fixation or extinction of the ancestral allele. This evolutionary process is modeled with a Continuous Time Markov Chain (CTMC) that jumps between two states at each locus.

First, we derive an expression for the expected state of the locus at time *t*. Given that the state of the locus at any point in time is a Bernoulli random variable with the success probability corresponding to the probability of the alternative allele being fixed, we can write the expected value for the state of locus *j* at time *t* as:

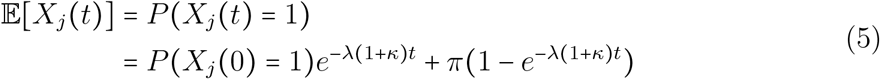

Second, we derive an expression for the the covariance in the state of the locus between two descendant lineages that diverged at time *t* and then evolved completely independently for *s* and *r* time units, respectively. This idea is illustrated in Figure 4(a). Here, we use the standard definition of the covariance:

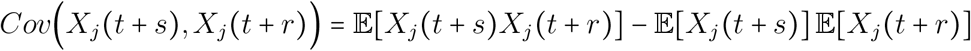

The unknown quantity in this case is the joint expectation of the state of the locus 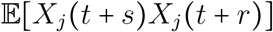, which we write in terms of the joint probability that the locus is fixed for the alternative allele in both lineages:

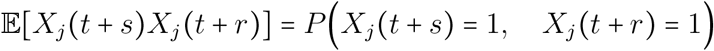

In order to express this in quantities that we have previously defined, we make use of the Law of Total Probability and factor the joint probability in the equation above into the marginal probability of the state of the locus in the ancestral population at time *t* and the probability of transitions in times *s* and *r* (see Figure 4(a)):

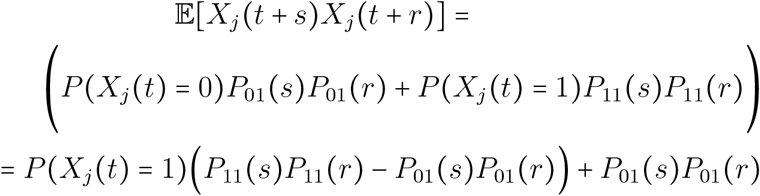

The factorization above assumes that the evolutionary process is independent after time *t*. As we will discuss later, this quantity will be most relevant in computing phylogenetic covariance of species that split at time *t* and then evolve for *s* and *r* time units completely independently. However, it is general and will be used to compute expressions for the time-dependent variance when we set *r* = *s* = 0. After substituting and simplifying, we obtain the following expression for the phylogenetic covariance in the state of the locus in the two descendant lineages:

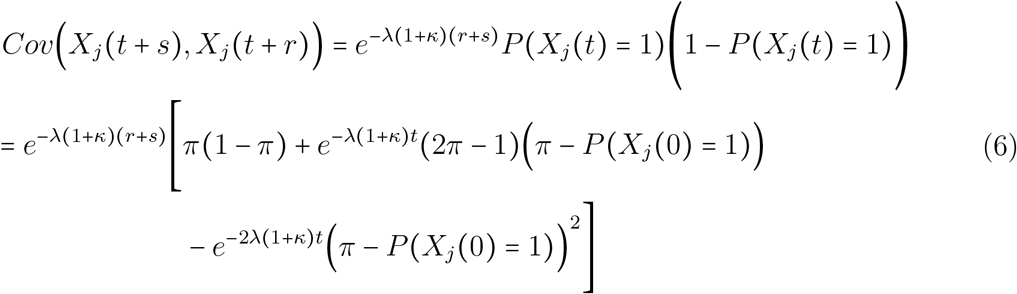

The above expression indicates that as s and r become large, the phylogenetic covariance for any given locus *j* decays exponentially towards zero.

### 1.2 Mean phenotype statistics

We defined the mean phenotype as a linear function of L bi-allelic loci whose state at any point in time we model with independent CTMCs with 2 states, corresponding to the reference and alternative alleles. The equation for the mean phenotype in a given population is:

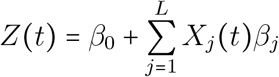

In this section, we derive expressions for the moments of this mean phenotype as a function of time and in the process we will comment on the dynamics that we observe.

#### 1.2.1 Phylogenetic mean of mean phenotype

In order to obtain expressions for the phylogenetic or ensemble mean of mean phenotype, we take the expectation with respect to all the random variables:

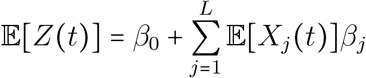

The above equation follows from the linearity the expectation and the assumption of independence between sites. If we plug in the definition of 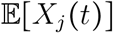 from Equation 5 into the equation above, we obtain:

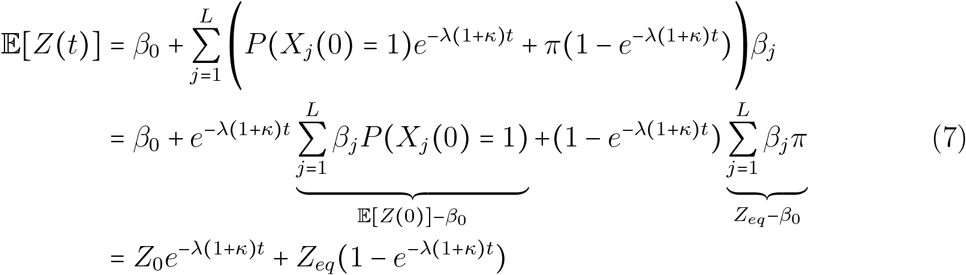

Where 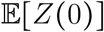 is the expected value of the mean phenotype at *t* = 0 and 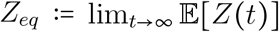 is the expected value of the mean phenotype at equilibrium. In phylogenetic modeling, we often assume that there is a single founding population at time *t* = 0, and we use this assumption to justify dropping the expectation from 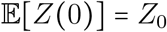. If we were to fit this model to data, both the initial phenotype *Z*_0_ and the equilibrium mean phenotype *Z_eq_* would be free parameters.

**Figure 3:**
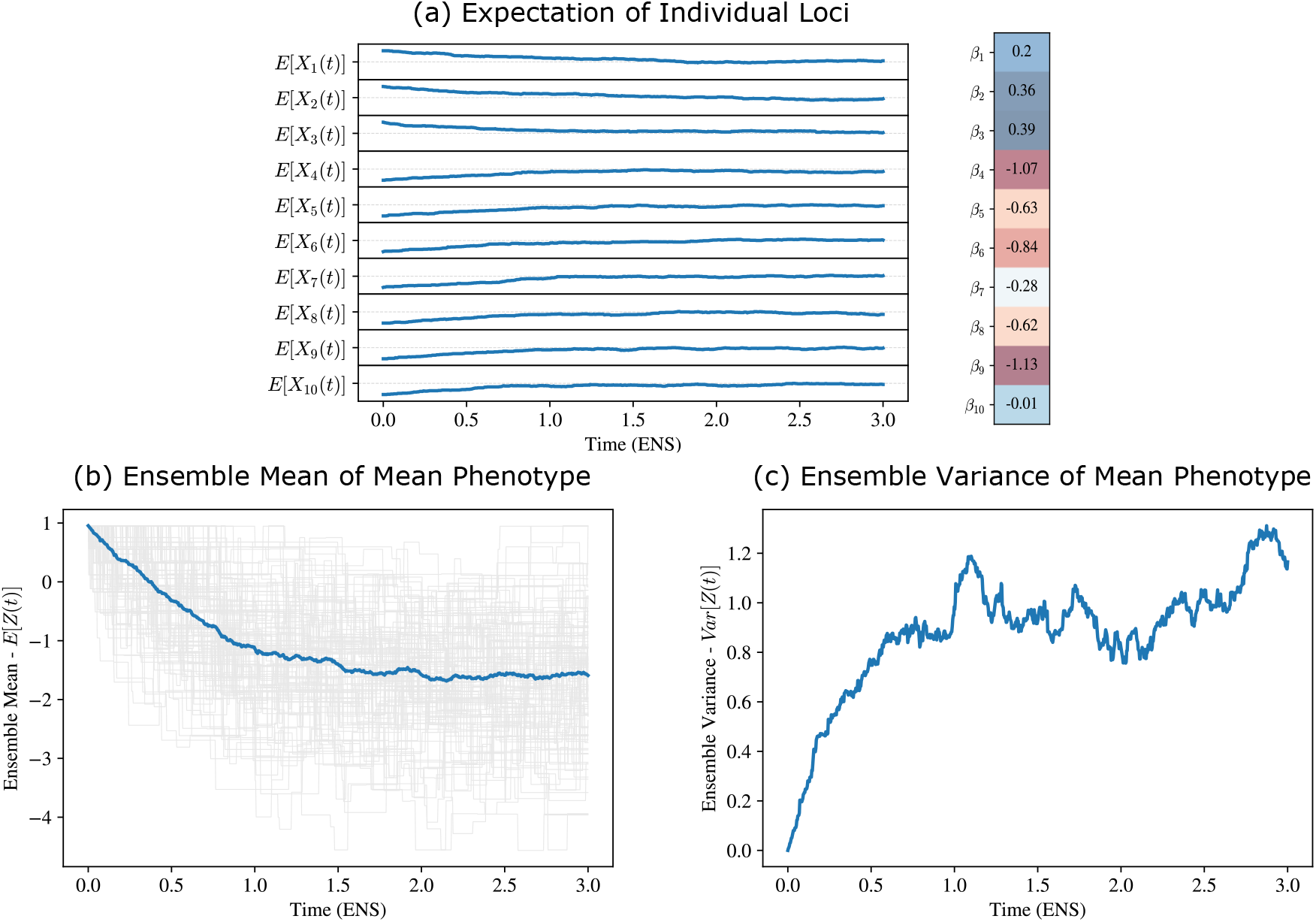
The ensemble mean and variance in mean phenotype as a function of time. Panel **(a)** shows the single locus statistics for *L* = 10 bi-allelic loci, which are obtained by averaging over the state of each locus in *N* = 100 independent replicates. In the ancestral population, the first three loci were fixed for the alternative allele while the remaining loci were fixed for the reference allele. Given the effect sizes ***β***, this results in an ancestral mean phenotype of *Z*_0_ ≈ 1. In this simulation we set *κ* = 1 for all loci and thus at equilibrium the expected value tends towards *π* = 0.5 for each locus. Panel **(b)** shows the mean phenotype in those replicate populations as a function of time (gray lines) as well as the ensemble mean 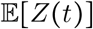 (blue line) that is obtained by averaging over the mean phenotype in all replicates. Panel **(c)** shows the empirical variance in mean phenotype across the different replicates. Note the variance saturates and reaches its equilibrium roughly twice as fast as the ensemble mean.

The above expression for the mean has the exact mean-reverting form as the mean of the Ornstein-Uhlenbeck process, as claimed in the main text and illustrated in Figure 3(b). Formulated in this way, we see that the role of mutation bias is to modulate the strength of the mutational force on the mean phenotype: if the bias is in the direction of the alternative allele, the mutational equilibrium is obtained faster, while if the bias is toward the reference allele, the equilibrium is obtained slower. In practice, since the bias would be on the order of 1, the effect on the mutational force is small, and with no bias, *κ* = 1, the force of mutation remains.

#### 1.2.2 Phylogenetic covariance of mean phenotype

The second quantity that we need to characterize the evolutionary dynamics of neutral phenotypes is the covariance, particularly the phylogenetic covariance between two or more species. The scenario that we envision in this case is that of an ancestral population that evolved for *t* time units and then split into two distinct species that evolved completely independently. The first species evolved for *s* time units and the second species evolved for *r* time units (Figure 4). This general characterization is flexible and allows for modeling non-ultrametric phylogenetic trees, which are common in studies of molecular evolution.

We use the definition of the covariance between two random variables and then plug in the definitions and simplify to obtain:

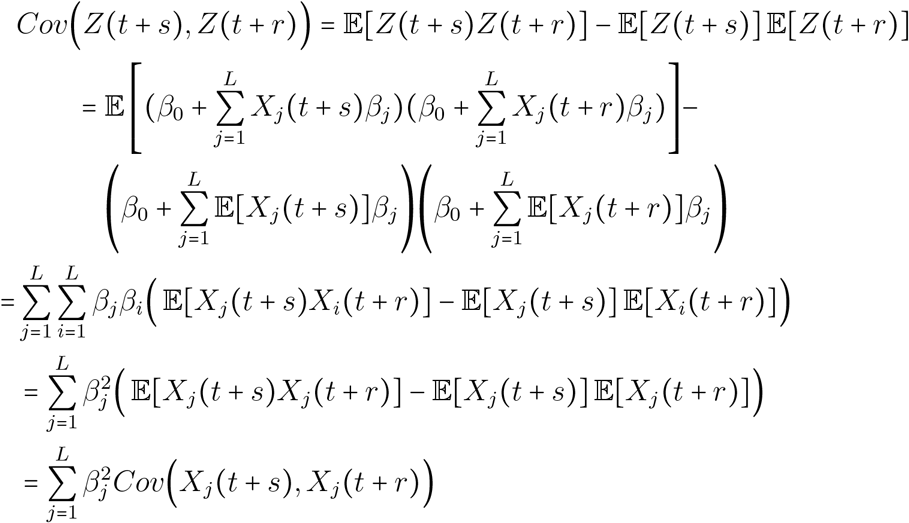

The last equality is due to the assumption of independence between sites. The phylogenetic covariance then reduces to the sum of the covariances between the state of each locus *j* at times (*t* + *s*) and (*t* + *r*), which we defined in the previous section. If we substitute the definitions and simplify, we obtain:

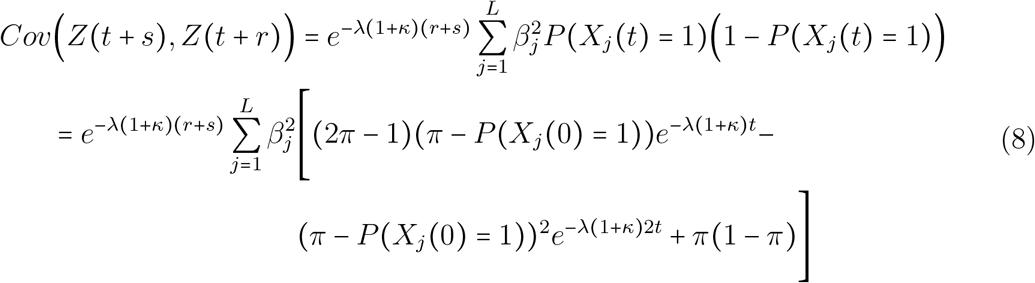

In this case, the covariance cannot be straightforwardly expressed in terms of phenotypic quantities such as *Z*_0_ and *Z_eq_*, and therefore depends on the genotype. In Section 3, we will show how to abstract away these genotypic quantities in order to perform inference at the phenotypic level.

**Figure 4:**
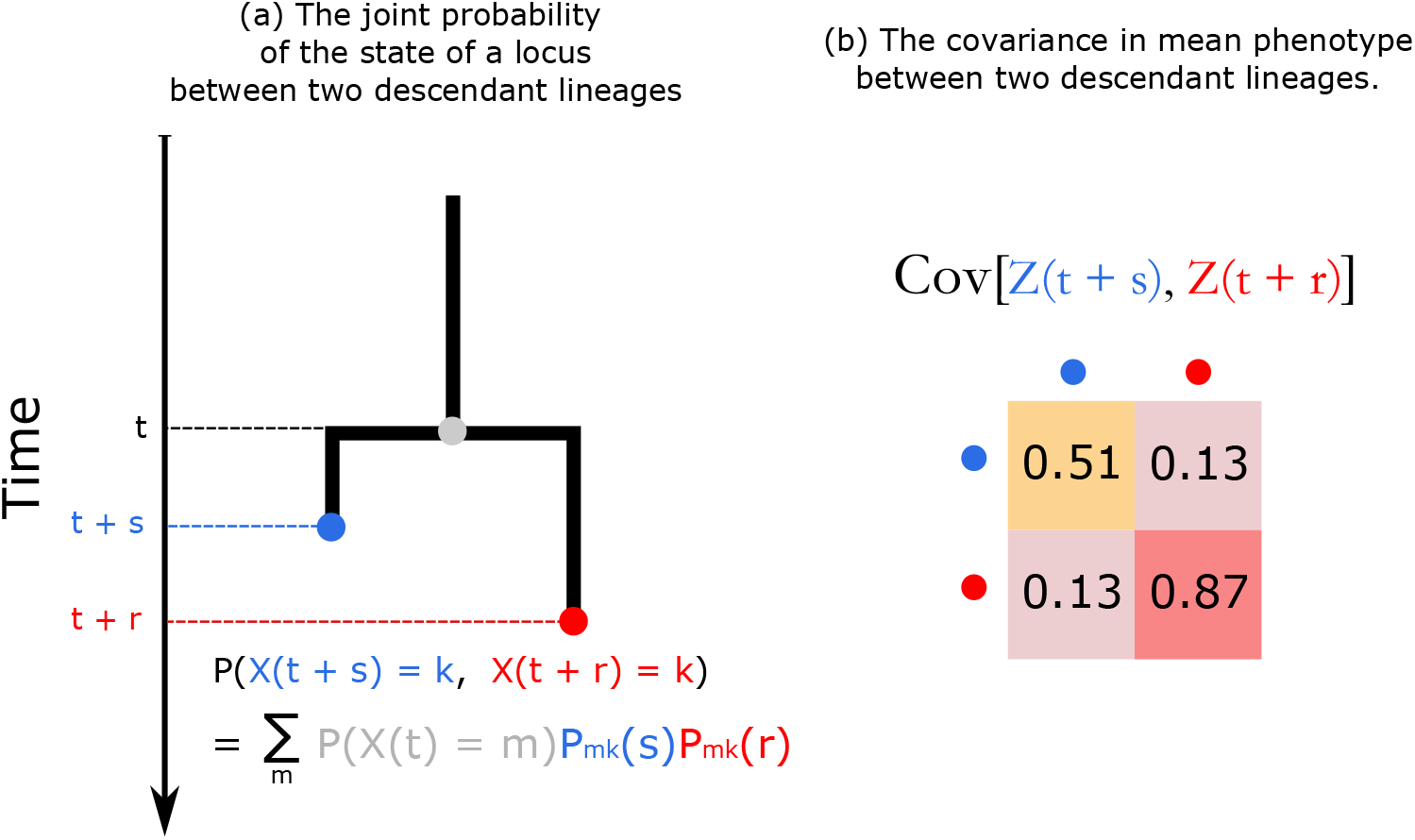
The phylogenetic covariance in mean phenotype as a function of time. In panel **(a)**, we illustrate the decomposition of the joint probability of the state of a locus *X_j_* in two descendant lineages at times *t* + *s* and *t* + *r* into the marginal probability of the state of the locus in the ancestral population at time *t* and the probability of *independent* transitions in times *s* and *r*. Panel **(b)** shows a numerical example of the covariance matrix of the mean phenotype between those two descendant lineages.

#### 1.2.3 Phylogenetic variance of mean phenotype

We can use the time-dependent covariance equation that we derived in the previous section to provide an explicit form for the phylogenetic variance of mean phenotype as a function of time. By substituting *r* = *s* = 0 and simplifying, we obtain:

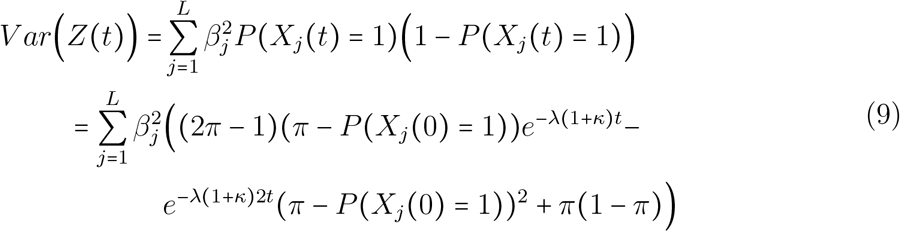

Unlike what the Brownian Motion model of phenotypic evolution predicts, this expression for the variance implies that it tends towards a stable equilibrium in the limit as *t* → ∞ (see Figure 3(c)):

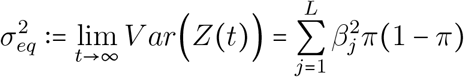

Where we define 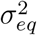 as the ensemble equilibrium variance of mean phenotype.

## 2 K alleles

While the bi-allelic model is useful for the purposes of deriving insights into the evolutionary dynamics of simple traits, in reality many of the the traits that are of interest to molecular biologists are underpinned by multi-allelic genetic architectures. This includes DNA sequence properties, such as GC content, as well as biophysical properties of peptides,such as net charge of a protein. When examining these types of molecular phenotypes on phylogenetic time-scales, accounting for multiple alleles becomes important. In this scenario, we model the phenotype as a linear function of *L* loci, each of which can have up to *K* different alleles:

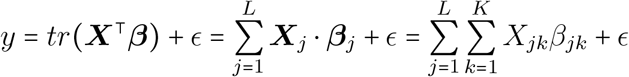

Where *tr*() is the trace operator, ***X*** is a *L* × *K* genotype matrix with each column indicating which of the *K* alleles is fixed, and ***β*** is the corresponding matrix of effect sizes. In this setup, *X_jk_* is a binary random variable indicating whether the *k^th^* allele is fixed at the *j^th^* locus. One main deviation from the bi-allelic model outlined previously is that we no longer maintain an intercept or a “reference allele”, instead keeping track of all alleles simultaneously.

We make use of the same assumptions that we employed in deriving the dynamics of the bi-allelic model, giving the following expression for the mean phenotype in a single population as a function of time:

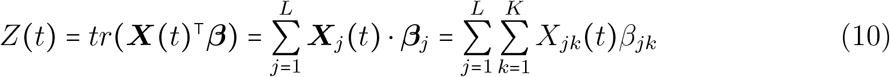

### 2.1 Single locus statistics

In the multi-allelic case, we model the genotype at each locus *j* with a CTMC that jumps between *K* discrete states, corresponding to the *K* different alleles. For simplicity, we make use of a generator matrix that is similar to that of the Felsenstein (1981) model of DNA evolution [5, 14]. This model gives the following transition probabilities for a given allele *k*:

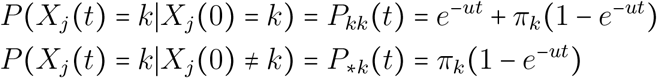

Here, *X_j_*(*t*) = *k* denotes that the *k^th^* allele is fixed at the *j^th^* locus at time *t* and *π_k_* is the equilibrium probability of observing the *k^th^* allele. The first equation is the conditional probability of the *k^th^* allele staying fixed after a time interval *t*, given that it was fixed in the founding population. The second equation gives the probability of transitioning to the *k^th^* allele after a time interval of *t*, given that any of the other alleles was fixed in the founding population. Unlike in the bi-allelic model, here we measure time in units of expected number of substitutions per site and thus define the scaling parameter *u* as:

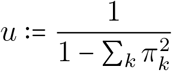

The equations above provide the probability of fixation or extinction in a given time interval conditional on the ancestral state. To get the probability that the *k^th^* allele is fixed at any point in time, we marginalize over the ancestral state:

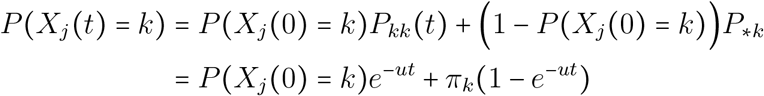

With this in hand, we can derive the single locus statistics that we would need to provide expressions for the dynamics of the mean phenotype. First, the expected value for the quantity *X_jk_*(*t*) becomes:

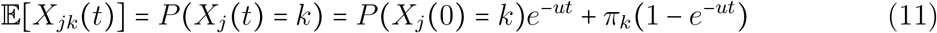

In addition to this, we need expressions for the time-dependent covariance between the allelic state of the locus j in two descendant lineages (see Figure 4). However, unlike the bi-allelic model, here we have to consider 2 scenarios: *Cov*(*X_jk_*(*t* + *s*), *X_jk_*(*t* + *r*)) and *Cov*(*X_jk_*(*t* + *s*), *X_jm_*(*t* + *r*)). The first scenario corresponds to the covariance in the state of the allele (fixed vs. extinct) in the two lineages, while the second scenario captures the covariance between the state of 2 different alleles (indexed by *k* and *m*). To derive expressions for those covariances, we first start by writing out expressions for the joint expectation. As we did previously in deriving the dynamics of the bi-allelic model, we make use of the Law of Total Probability to decompose the joint probabilities into familiar quantities.

For the first covariance term, we decompose the joint probability into the marginal probability of the state of the locus in the ancestral population at time *t* and the probability of independent transitions to the *k^th^* allele in times *s* and *r*:

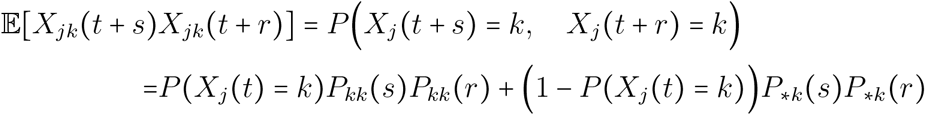

The decomposition for the second covariance term is similar. However, in the ancestral population at time *t* there are three possible cases, mainly that the locus was fixed for the *k^th^* allele, for the *m^th^* allele, and for neither allele:

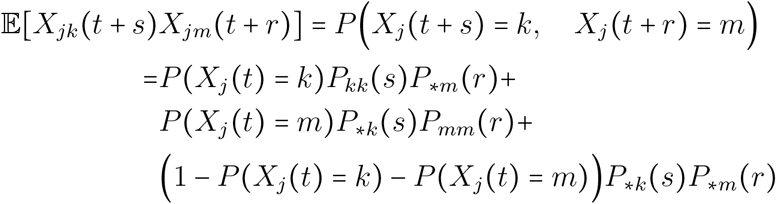

With this in hand, after plugging in the definitions and simplifying the algebra, we can write the single-locus covariances as:

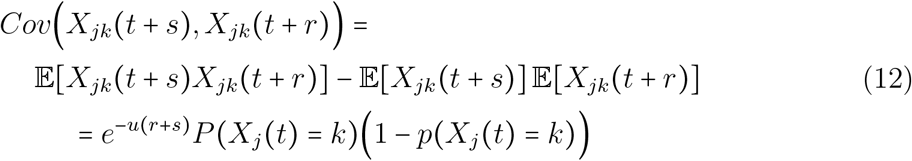

Which we can expand to obtain:

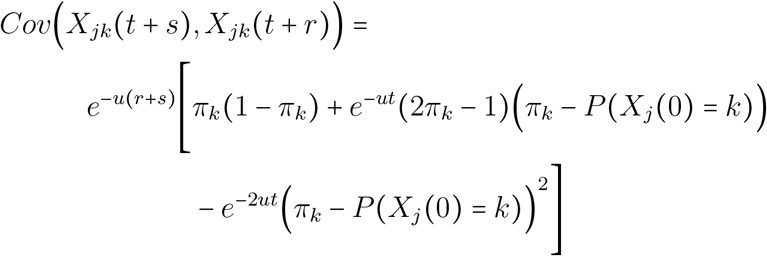

And the covariance of two different alleles can be written as:

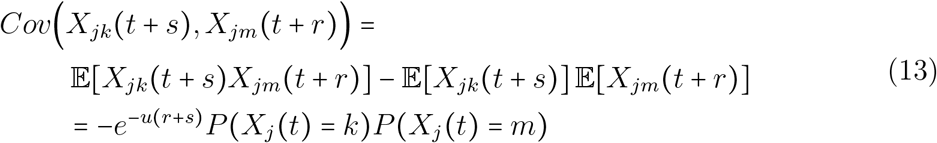

Naturally, the first covariance term resembles the expression for the single locus covariance that we derived for the bi-allelic model. The second covariance term is unique, since it captures the covariance between 2 distinct alleles. With these quantities in hand, and remembering that we still maintain the assumption of independence between sites, we can derive the mean phenotype statistics, which we turn to in the next section.

### 2.2 Mean phenotype statistics

Recall that we defined the mean phenotype as a linear combination of L loci whose state at any point in time we model with independent CTMCs with *K* states corresponding to the *K* distinct alleles. The equation for the mean phenotype in a given population is given by:

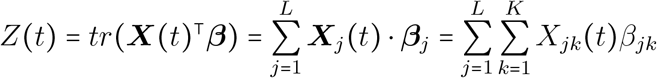

#### 2.2.1 Phylogenetic mean of mean phenotype

In order to obtain expressions for the phylogenetic or ensemble mean of the mean phenotype, we take the expectation with respect all random components of this model:

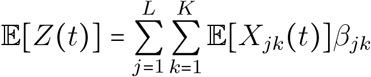

The above equation follows from the linearity the expectation and the assumption of independence between sites. If we plug in the definition of 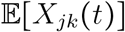 from above, we obtain:

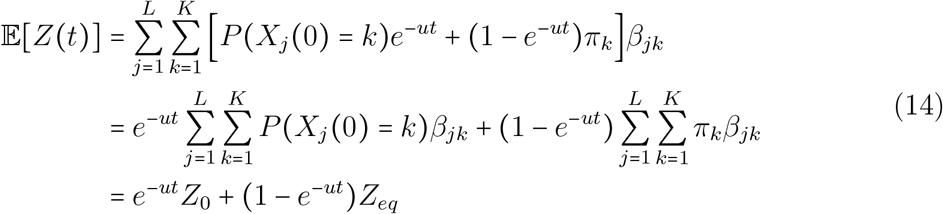

Where we define 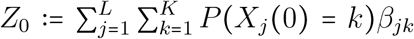 and 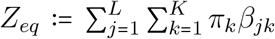 as the mean phenotype in the founding population and the ensemble mean at equilibrium, respectively. Similar to what we saw with the bi-allelic model, the expresion for the mean has the exact form of the mean dynamics of the Ornstein-Uhlenbeck process.

#### 2.2.2 Phylogenetic covariance of mean phenotype

To derive an expression for the covariance of the multi-allelic model, we use the definition of the covariance and simplify:

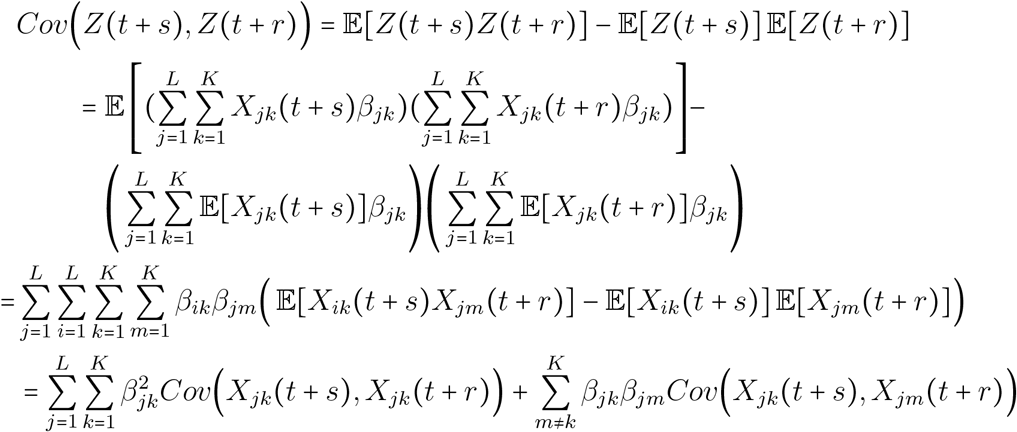

The last equality holds because of the assumption of independence between sites. After plugging in the single-locus covariances that we derived previously, we can write the phylogenetic covariance in mean phenotype as:

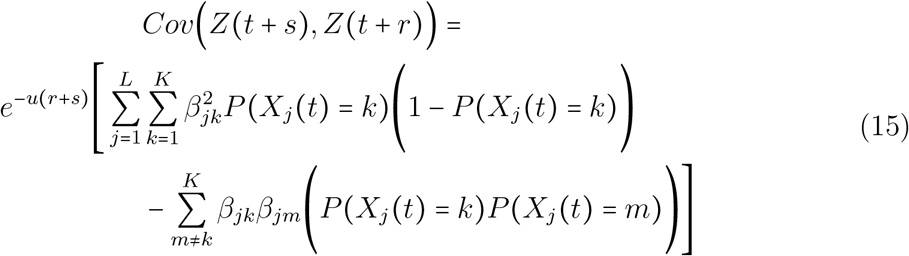

#### 2.2.3 Phylogenetic variance of mean phenotype

To obtain an expression for the phylogenetic variance of mean phenotype as a function of time, we simply set *s* = *r* = 0 in the covariance equation we obtained above, giving:

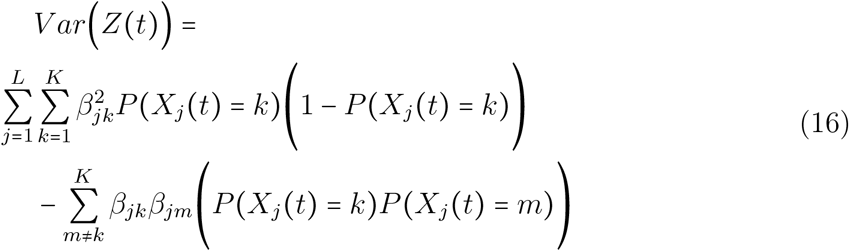

We note that the expression for the variance corresponds to the variance of a categorical random variable with *K* outcomes, as expected. In general, the variance appears to depend on the initial genotype and is non-monotonic. Furthermore, in line with the results from the bi-allelic model, the variance in this model is bounded. The ensemble equilibrium variance can be obtained by taking the limit as *t* → ∞, giving:

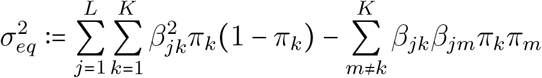

## 3 Simplifying and extending the model

### 3.1 Lumping alleles and the grouped model

In this section, we will show that the multi-allelic models can be vastly simplified for a wide-range of traits by only tracking the state of a number of alleles that is equivalent to the *unique* number of effect sizes. For example, for counting-based traits such as GC Content, instead of modeling the contribution of all four DNA letters to the dynamics, it suffices to keep track of two “lumped” or grouped alleles, corresponding to the G/C and non-G/C state. This construction will help simplify analyzing the evolutionary dynamics of many molecular and cellular traits.

In particular, we will show that the ensemble mean and covariance dynamics under the grouped model are almost identical to the dynamics under the full model, with a an important difference in the scaling factor that we will comment on shortly. To illustrate, we assume that out of the *K* alleles that characterize the underlying sequence, an arbitrary subset, denoted by *S*, have an effect size of *β*_1_ while the remaining alleles have an effect size of *β*_2_:

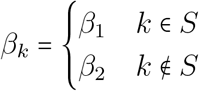

Furthermore, since the fixation of alleles in a given population constitutes disjoint events by construction, we can write the probability of observing any of the alleles in the subset *S* at time *t* as:

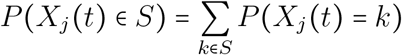

With the complement of this being *P*(*X_j_*(*t*) ∉ *S*) = 1 – *P*(*X_j_*(*t*) ∈ *S*). And because this is a bi-allelic model where the alleles are the states *S* and its complement, we observe that the single locus covariance can be written as:

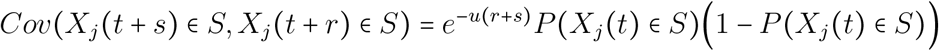

#### Phylogenetic Mean of Mean Phenotype

Given our construction above, we can write the contribution of a single locus *j* to the phylogenetic mean under the grouped model as:

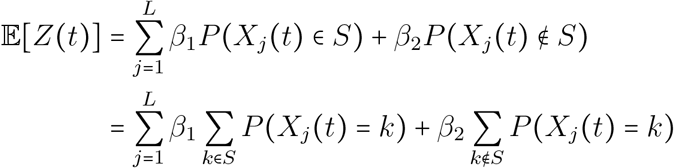

Thus, the dynamics of the phylogenetic mean under the full and grouped models are identical. In the case of the mean, it is straightforward to generalize this to any arbitrary grouping of alleles by their effect size.

#### Phylogenetic Covariance of Mean Phenotype

Showing that the equivalence holds for the covariance requires more work. In order to show this, it helps to re-write the covariance in the full model as follows and then simplify:

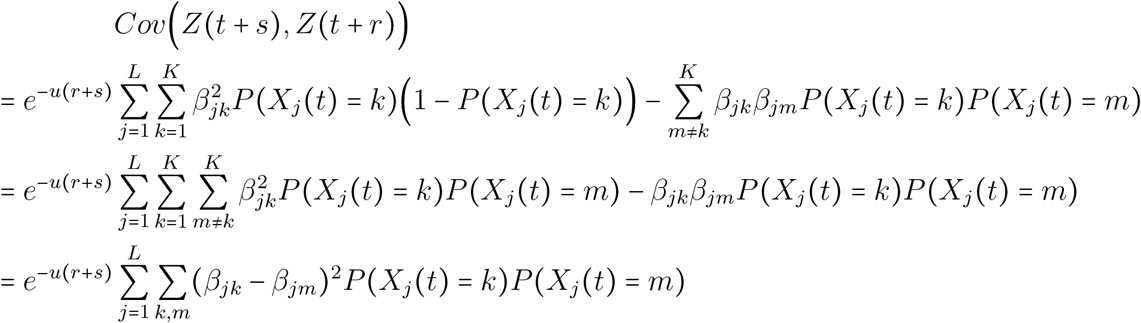

Where the sum Σ_*k,m*_ is over *unique pairs* of alleles *j* and *m*. The second equality follows from the fact that *P*(*X_j_*(*t*) = *k*)(1 – *P*(*X_j_*(*t*) = *k*)) = Σ_*m≠k*_ *P*(*X_j_*(*t*) = *k*)*P*(*X_j_*(*t*) = *m*). When an effect size is shared between a pair of alleles, then (*β_jk_* – *β_jm_*)^2^ = 0 and their joint contribution to the covariance drops out of the sum. If there are only two unique effect sizes *β*_1_ and *β*_2_, then the sum above would simplify to:

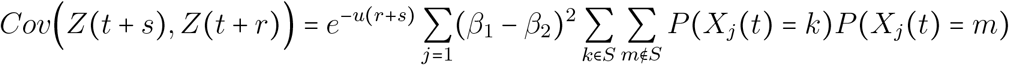

Where *S* is the subset of alleles with effect size *β*_1_. This expression is exactly what we would obtain if we solved for the covariance in the grouped bi-allelic model and then plugged in the definitions of the probabilities from before:

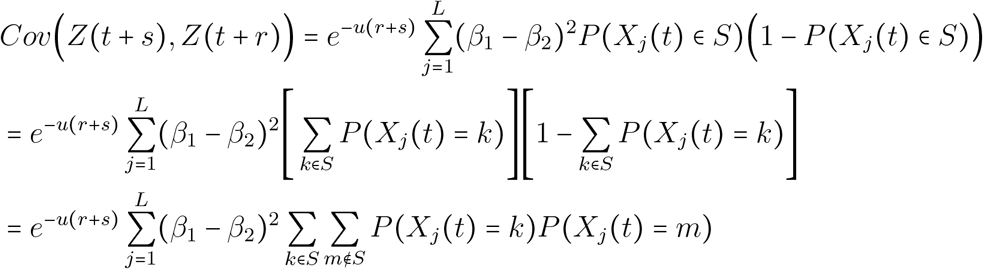

Therefore, we have shown that the covariance dynamics under the full and grouped models are identical.

#### The scaling factor *u* under the grouped model

An important detail that we omitted in the preceding discussion is the scaling factor *u* and whether it is the same or takes on a different value under the grouped model. A naive interpretation may suggest the *u* under the grouped model should be defined as:

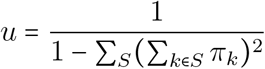

Where the sum in the denominator is over the grouped alleles instead of the individual alleles in the underlying sequence. However, this is the major difference between the full model and the grouped model: the scaling factor has to be defined in terms of the equilibrium probabilities of all alleles:

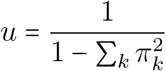

To see that this is the case, recall that in our construction of this grouped formulation, we defined the probabilities of observing the grouped allele as a sum of the probabilities of observing the individual alleles. And in these individual probabilities, the scaling factor is defined under the full model. Therefore, under the grouped model, u retains the same value and interpretation as under the full model.

### 3.2 Abstracting away the genetic architecture from the covariance

In our analysis of the model, we noted that the dynamics of the mean can be expressed in phenotypic quantities, abstracting away the underlying genetic architecture of the trait, including the number of causal loci and their effect sizes. Here, we will show that a similar abstraction can be achieved in the case of the covariance.

The covariance involves many terms that are harder to abstract away. This is mainly the case because of the cross-terms that describe the covariance between different alleles at each site. However, it is possible to write the covariance in terms that completely abstract away from the underlying genetic architecture. To see this, we start with the formulation of the covariance that we derived in the description of the grouped model (Section 3.1):

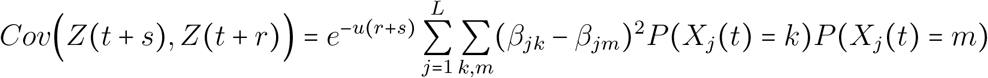

Recalling the fact that *P*(*X_j_*(*t*) = *k*) = *P*(*X_j_*(0) = *k*)*e^−ut^* + *π_k_*(1 – *e^−ut^*) and the assumption that we have a single ancestral monomorphic population (thus *P*(*X_j_*(0) = *m*)*P*(*X_j_*(0) = *k*) = 0), it is straightforward to show that the above expression simplifies to:

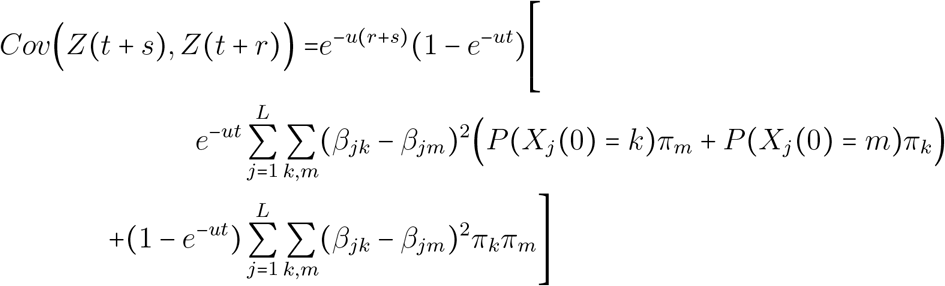

This expression can be further simplified by observing that when we set *r* = *s* = 0 and take the limit as *t* → ∞, we get that the sum on the third line equals the ensemble equilibrium variance that we have previously defined: 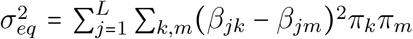. Furthermore, introducing a new quantity that we denote by Ψ, we define:

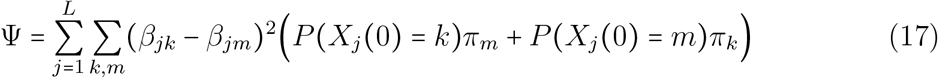

The quantity Ψ does not have a straightforward interpretation in general, though it is always positive and we may gain insights into its role by analogy with the Gaussian models of trait evolution. For instance, if the evolutionary process starts at equilibrium (i.e. *P*(*X_j_*(0) = *k*) = *π_k_*), then we see that Ψ takes on the value of 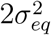. By comparison, under the Ornstein-Uhlenbeck model, the equilibrium variance is 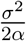 [1]. Furthermore, as we will show in Section 3.4, the quantity *u*Ψ determines the rate at which the variance increases at small time. Under the OU model, on the other hand, this behavior is controlled by the parameter *σ*^2^. All of this is suggestive of the fact that the quantity *u*Ψ acts in similar ways to behavior of the dispersion parameter *σ*^2^ in the Ornstein-Uhlenbeck model.

With this in hand, we can write the covariance as:

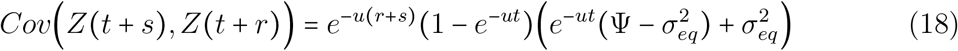

Thus, we have arrived at an expression for the covariance in mean phenotype between two descendant lineages that abstracts away the underlying genetic architecture.

### 3.3 Phylogenetic divergence of mean phenotype

The expressions for phylogenetic mean and covariance derived in previous sections are sufficient to construct a likelihood-based inference framework to fit our model to data. Indeed, these quantities have been the central focus of analysis and extensions in the Phylogenetic Comparative Methods community [3, 7]. However, a more recent theoretical framework inspired by techniques from statistical physics has shown that another useful quantity to differentiate between the dynamics of neutral and adaptive evolution is the phylogenetic divergence of mean phenotype [11, 8, 10]. Defined as the expected squared difference in the mean phenotype in two extant lineages, the divergence can be expressed in terms of quantities that we defined above. Mathematically, we can write the pairwise divergence between the mean phenotypes of the two lineages as:

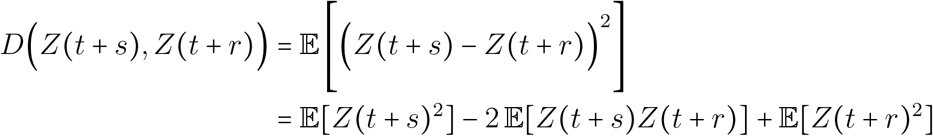

By observing that 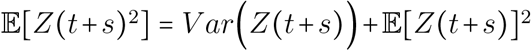 and that 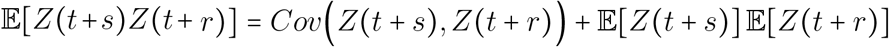, we can see that the phylogenetic divergence can be expressed in terms of the mean, variance, and covariance in the mean phenotype between the two lineages:

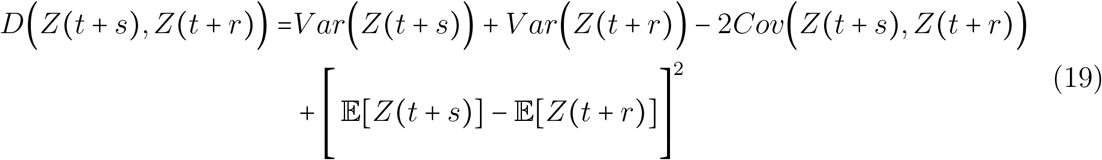

This is a general expression that can work with any model of phenotypic evolution. Under our model, the expression can be a bit unwieldy, though by examining the dynamics of this expression at equilibrium, we recover some of the general formulations for quantities that were defined in the framework of Nourmohammad et al. [11, 10]:

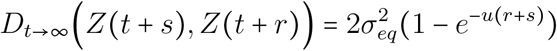

In the limit of long divergence times (as *r* + *s* becomes large), we see that this quantity becomes twice the equilibrium variance 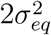, which in this context is the *multi-allelic* generalization of the trait scale *D*_0_ as defined in [11, 10].

### 3.4 Linear approximation for the variance at small time

In this section, our aim is to examine the dynamics of the ensemble variance at small time (*ut* ≪ 1) and to obtain a linear approximation in this regime. Starting with the compact representation of the covariance in Equation 18 and setting *r* = *s* = 0, we obtain the following expression for the ensemble variance at time *t*:

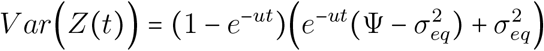

At small time, we have that 1 – *e^−ut^* ≈ *ut* and (1 – *e^−ut^*)*e^−ut^* ≈ *ut*, which gives the following linear expression for the variance at small time:

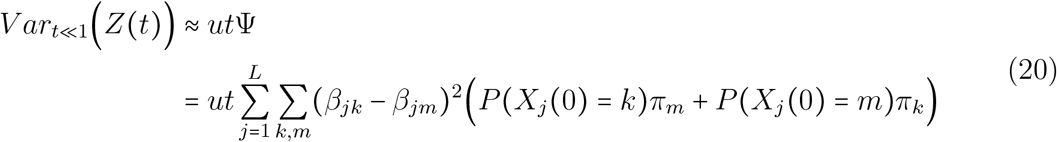

Thus, the variance dynamics at small time only depend on the quantity *u*Ψ, which we defined in previous sections and shown to be related to the dispersion of the evolutionary process.

### 3.5 Simplifying covariance in the bi-allelic model

There are a number of simplifying assumptions that we can invoke to gain insights into the behavior of the time-dependent covariance in the bi-allelic model.

#### Covariance at equilibrium

One common assumption states that the evolutionary process starts in an equilibrium state and thus *P*(*X_j_*(0) = 1) = *π*. This simplifies the expression for the covariance in the bi-allelic model to:

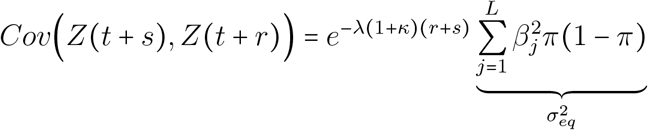

Where 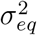 is the equilibrium variance of the mean phenotype *Z*. In this case, 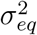 would be a free parameter in the model that can be fit to data. While the form of this is attractive, it is inconsistent with the weak-mutation assumption and empirical observations that genotypes are not at their equilibrium in finite populations, but rather close to monomorphic. For example, this formula predicts that the cross-species variance is always at its mutational equilibrium.

#### Covariance with no mutational bias

A different simplification is obtained if we assume that the mutation rate is symmetric between the 2 alleles (i.e. *κ* = 1) and that there was a single founding population (i.e. *P*(*X_j_*(0) = 1) ∈ {0,1}). The latter assumption is common in phylogenetic analysis and it implies that the initial variance is zero. Under this scenario, we obtain that *π* = 0.5, and therefore the covariance simplifies to:

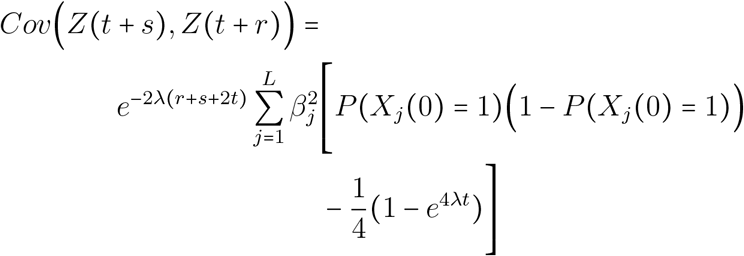

By observing that *P*(*X_j_*(0) = 1)(1 – *P*(*X_j_*(0) = 1)) = 0 and recalling the definition of the equilibrium variance, we can write the covariance under these assumptions as:

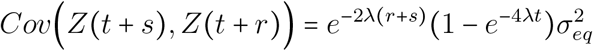

By setting *s* = *r* = 0, we also obtain an expression for the variance under this model, which turns out to match exactly the variance dynamics of the Ornstein-Uhlenbeck process, as alluded to previously:

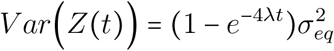

**Figure 5:**
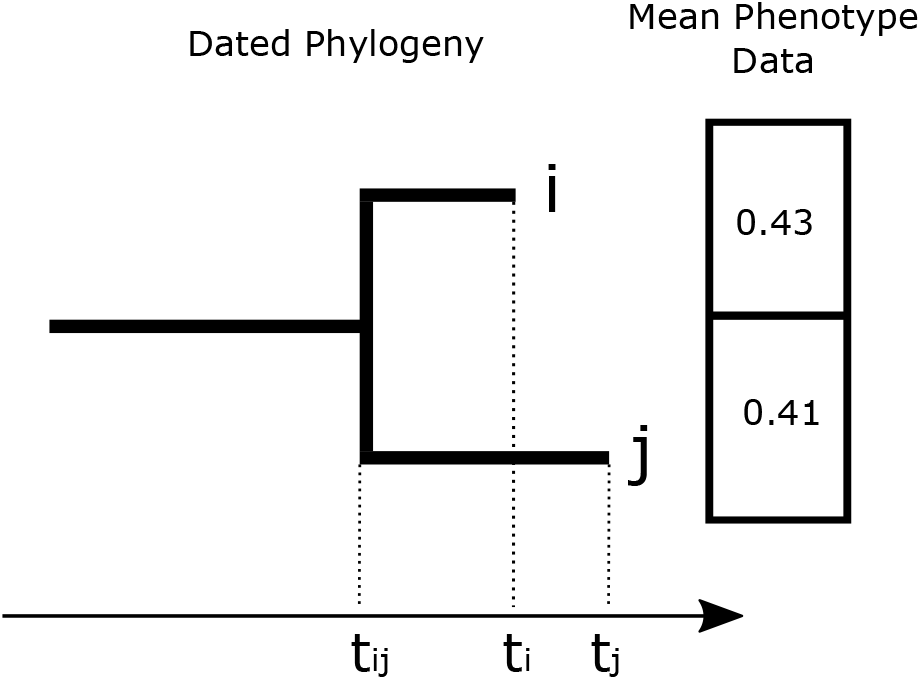
An illustration of the phylogenetic inference setup. In typical applications, we have a dated phylogeny, where time is in units of Expected Number of Substitutions (ENS), as well as mean phenotype measurement in extant species. These inputs are typically combined with a parametric model, e.g. a multivariate Gaussian likelihood, whose parameters are optimized using Maximum Likelihood.

## 4 Phylogenetic inference under the model

The theoretical model that we outlined in the previous sections provides a general picture of the dynamics of the moments of neutral phenotypes on phylogenetic time-scales. A natural extension of this model would be to use it for phylogenetic inference tasks that are popularly encountered within the realm of Phylogenetic Comparative Methods (PCMs). For example, this model can be used to reconstruct the ancestral mean phenotype, assuming that we have sufficient evidence that the phenotype evolves under little selective constraint. Another important application of this model is to provide a more powerful test for selection, replacing Brownian motion as the null model of phenotypic evolution. For these and other tasks, we need to derive a parametric model for the evolutionary dynamics of mean phenotype that can be fit to data using a Maximum Likelihood framework.

### 4.1 The likelihood

We assume that we are given a (rooted) phylogeny that describes the relationships between *N* species {*S*_1_,…, *S_N_*} and the divergence times that separate them in units of Expected Number of Substitutions per site (ENS). In this setup, we assume that species *i* is removed from the root of the phylogeny by *t_i_* time units and that a pair of species *i* and *j* diverged at time *t_ij_* and then each evolved for *t_i_* – *t_ij_* and *t_j_* – *t_ij_* time units, respectively. Additionally, assuming that we have measurements of the mean phenotype in each of those species ***Y*** = {*y*_1_,…, *y_N_*}, our aim in this section is to define a likelihood for the data given the parameters of the model 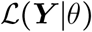. See Figure 5 for an illustration of this setup.

Our theoretical model does not describe an explicit probability distribution. However, assuming that the trait of interest is sufficiently Gaussian, we can approximate its time-dependent dynamics with a Gaussian Process that is characterized by the following mean and covariance terms:

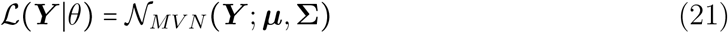

Where 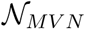 is a multivariate Gaussian distribution with *N* coordinates and where the entry in the mean vector for species *i* is defined as:

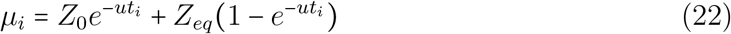

And the covariance between any pair of species *i* and *j* is defined as:

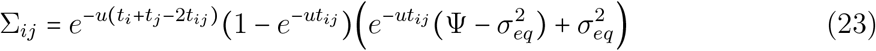

This most general model has a total of 5 parameters: 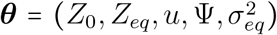. However, note that for many traits and species of interest, the parameter *u* can be fixed to reasonable values based on genome-wide estimates of the equilibrium probabilities of the 4 DNA letters. For instance, assuming a Jukes-Cantor (1969) model of DNA evolution [9, 14], *u* may be preset to 1.3. This results in a neutral null model of phenotypic evolution with 4 parameters to estimate, similar to the standard formulation of the phylogenetic Ornstein-Uhlenbeck model [7]. In some applications, when analyzing simple linear traits, e.g. GC Content, other parameters such as 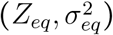 may also be fixed to reasonable values that can be straightforwardly derived based on the equilibrium probabilities of the 4 DNA letters. An example of this will be shown in Section 5.

While the general model has a relatively large number of parameters, these parameters are easily interpretable and it is straightforward for practitioners to examine their meaning or add reasonable constraints on their values in model fitting procedures. For example, the parameters *u* can be restricted to the range (1,2). Additionally, the parameters Ψ and 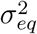 may be restricted to positive real numbers.

Finally, note that, like the Ornstein-Uhlenbeck model of phenotypic evolution, the covariance matrix predicted by our neutral model has a Generalized 3-Point Structure as identified by [13], making it amenable linear time model fitting procedures.

### 4.2 Comparison with Brownian motion and the Ornstein-Uhlenbeck process

To compare our model to the standard Gaussian models of phenotypic evolution, we first write out their likelihood.

#### Brownian motion

Under the Brownian motion model, the mean and covariance of the multivariate Gaussian distribution are given by [7]:

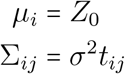

The Brownian motion model has only two parameters, *Z*_0_ and *σ*^2^, and is thus the simplest Gaussian model for phenotypic evolution. However, it predicts that the phenotypic variance increases linearly with time and does not saturate, violating intuitions and observations. Note that the Brownian motion model also predicts a linear, non-saturating pairwise divergence, as defined in Section 3.3 and in the framework of Nourmohammad et al. [11]. The pairwise divergence according to the Brownian motion model is given by:

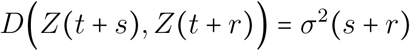

This disagrees with our model predictions as well as with the neutral null model of Nourmohammad et al. [11, 8, 10]. It is well known, however, that the Brownian motion model is a good approximation for the dynamics at small time *t*.

#### Ornstein-Uhlenbeck (OU) model

The OU model of phenotypic evolution, which is used to model the action of stabilizing selection, defines a Gaussian process likelihood for the data where the mean and covariance are given by [7]:

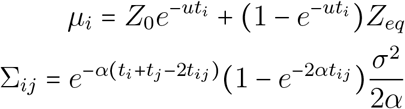

Similar to our model with fixed scale *u*, the OU model has a total of 4 parameters ***θ*** = (*Z*_0_, *Z_eq_*, *σ*^2^, *α*). However, in standard implementations, as in the `R` packages `ouch` [1] and `geiger` [12], practitioners assume that the ancestral phenotype is the same as the equilibrium phenotype (*Z*_0_ = *Z_eq_*), thus reducing the number of parameters to 3. As noted elsewhere in the Appendix and the main text, our model shares the exact form of the mean with the OU model, though the two models diverge in the form of the covariance.

At equilibrium, the OU model predicts a constrained pairwise divergence that is modulated by the parameter α, as we would intuitively expect and consistent with the action of the parameter c in the models of Nourmohammad et al. [11, 8, 10]:

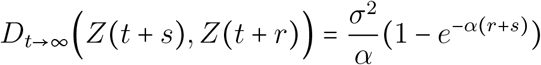

However, note that in standard applications of the OU model, the parameter *α* is often difficult to estimate accurately and there are debates about its meaning and interpretation in this context [2].

## 5 Examples and applications

In this section, we provide three examples that illustrate the utility and insights that can be derived from this modeling approach. In the first example, we derive explicit expressions for the time-dependent dynamics of the phylogenetic mean and variance of GC Content under the Felsenstein (1981) [5] model of DNA evolution. In the second example, we examine these same dynamics under the Jukes-Cantor (1969) [9] model of DNA evolution. Finally, in the third example we show that this approach can tackle more complex genetic architectures by examining dynamics of the number of ATG codons in a DNA sequence.

### 5.1 Dynamics of GC content under the F-81 model

Under the modeling framework that we outlined, our genotype matrix ***X*** and the corresponding matrix of effect sizes ***β*** are of dimension *L* × 4 for a sequence of length *L*. For a trait like GC Content, each row of the matrix of effect sizes has the following form:

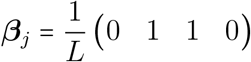

Under the F-81 substitution model, the four alleles have a stationary distribution specified by: ***π*** = (*π_A_, π_C_, π_G_, π_T_*). These stationary probabilities can be inferred from genome-wide data or from a set of sequences that are of interest to the analyst. Additionally, the scaling parameter *u*, which modulates the convergence rate to the equilibrium state, is given by [14]:

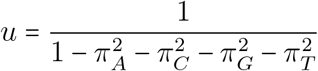

#### 5.1.1 Dynamics of the mean and variance

##### Dynamics of the Phylogenetic Mean

The dynamics of the mean for GC Content can be obtained straightforwardly from Equation 14 by plugging in the effect sizes:

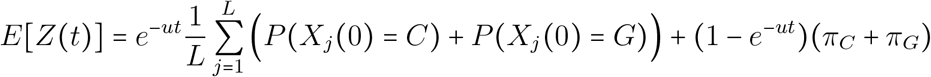

If we assume that there is a single founding population, then the quantity 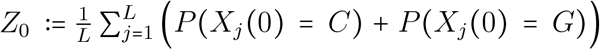 becomes simply the fraction of G/Cs in the ancestral sequence. In addition, as can be intuitively gleaned, the expected mean phenotype at equilibrium under this model is shown to be *Z_eq_*: = *π_C_* + *π_G_*. This example shows that the phylogenetic dynamics of the mean for some molecular traits can be explored analytically.

##### Dynamics of the Phylogenetic Variance

Similarly, we can obtain the dynamics of the variance for GC Content by plugging in the effect sizes into Equation 16 and simplifying:

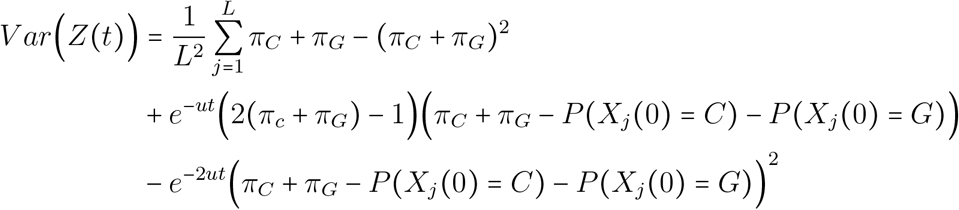

An interesting feature of the equation above is that it resembles the variance expression for the bi-allelic model, as if in effect we only have 2 alleles corresponding to the A/T and G/C states. This suggests that, in similar scenarios and for traits that have similar distributions of effect sizes, it makes sense to “lump” the alleles that share the same effect size into a single allele, thus reducing the number of states that we need to keep track of. Indeed, this supposition is shown to be true for any arbitrary grouping of alleles by their effect sizes in Section 3.1.

If we use the definitions of *Z*_0_ and *Z_eq_* from above, we can further simplify the expression for the variance as follows:

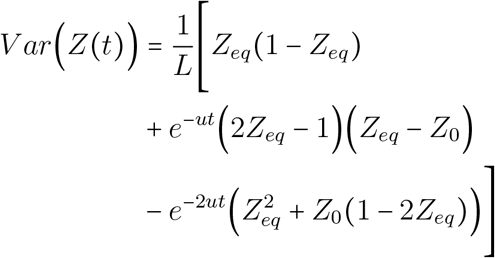

In this case, the phylogenetic variance is independent of the genotype. If we take the limit as *t* → ∞, we obtain the following expression for the equilibrium variance:

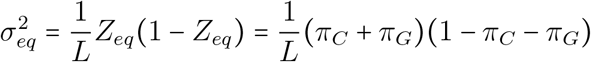

It is important to highlight here that the variance is inversely proportional to the number of loci *L*, as would be expected for traits that are expressed as a fraction of the length of the sequence.

#### 5.1.2 Linear approximation for the variance of GC Content under F-81

In the main text, we analyze the variance of GC Content at small phylogenetic timescales, on the order of the divergence of modern primates. Here, we derive the model predictions for the variance dynamics of GC Content in the regime when *ut* ≪ 1. In Section 3.4, we have shown that the variance at small time only depends on the parameter Ψ (see Equation 20):

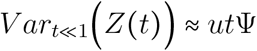

For GC Content, by noting that there are only 4 unique combinations of alleles that do not share effect sizes (*A/C, A/G,T/C, T/G*), it is straightforward to write Ψ in terms of other quantities, mainly *Z*_0_ and *Z_eq_*:

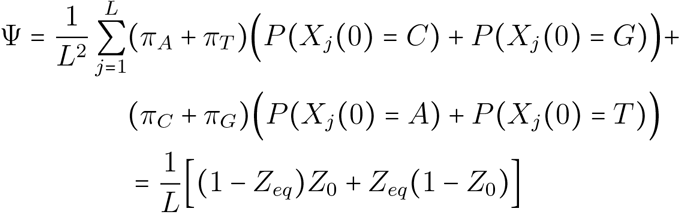

Which gives the following linear approximation for the variance of GC Content at small time:

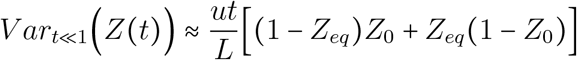

### 5.2 Dynamics of GC content under the JC-69 model

A special case of the example that we outlined in the previous section is obtained when we examine sequences evolving according to Jukes-Cantor model, which is specified by the following stationary distribution: 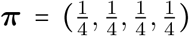. From this, we can see that the scaling parameter becomes 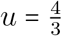.

#### 5.2.1 Dynamics of the mean and variance

##### Dynamics of the Phylogenetic Mean

Under this model, the phylogenetic mean has a simple form:

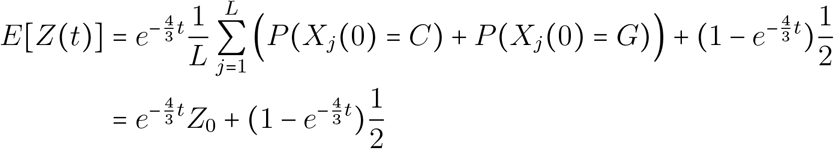

Where 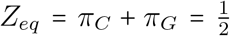 and *Z*_0_ again corresponds to the fraction of *G/C*s in the ancestral sequence.

##### Dynamics of the Phylogenetic Variance

To obtain an expression for the phylogenetic variance under the Jukes-Cantor model, we simply plug in 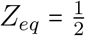 in the expression we derived for the F-81 model:

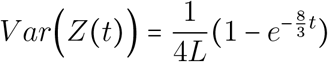

Because of the symmetry of the JC69 model, the expression above has the form of the variance as that of the Ornstein-Uhlenbeck model (i.e. a single exponential decay towards the equilibrium variance at a rate that is twice as fast as the rate of the mean reverting to its equilibrium value). The equilibrium variance under this model is given by:

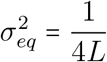

#### 5.2.2 Linear approximation for the variance of GC Content under JC-69

Using the expression we derived in section 5.1.2 and noting the 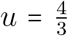 and 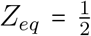, we can write the linear approximation for the variance dynamics at small time under the Jukes-Cantor model as:

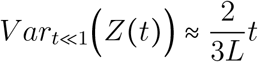

Similar to what we observed for the full-scale expression for the variance dynamics, the short timescale variance behavior is also independent of the initial phenotype *Z*_0_.

### 5.3 Dynamics of ATGs under the F-81 model

To showcase the generality of the framework we outlined above, here we analyze a more complex case involving codon counts in a DNA sequence. The trait of interest here is the number of *ATG* codons in a DNA sequence of length *L*. In this context, there are 64 alleles corresponding to the 64 distinct codons. Therefore, the scaling factor *u* is defined as:

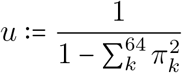

Where *π_k_* is the equilibrium probability of codon *k*, which can be derived from the equilibrium probabilities of the individual DNA letters. Since we are defining a countingbased trait, the effect sizes for all codons are 0 except for the *ATG* codon, which has an effect size of 1.

#### 5.3.1 Dynamics of the mean and variance

##### Dynamics of the Phylogenetic Mean

The dynamics of the mean for the number of ATGs can be obtained straightforwardly from Equation 14 by plugging in the effect sizes:

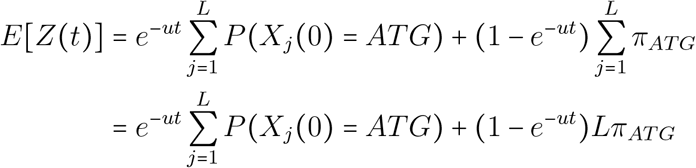

Assuming a single founder population, we can see that 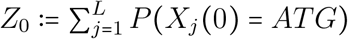 is the number of *ATG* codons in the ancestral sequence and *Z_eq_*: = *Lπ_ATG_* is the expected number of ATGs at equilibrium.

##### Dynamics of the Phylogenetic Variance

Similarly, we can obtain the dynamics of the variance for GC Content by plugging in the effect sizes into Equation 16 and simplifying (it may be useful to refer to the results in Section 3.1 here):

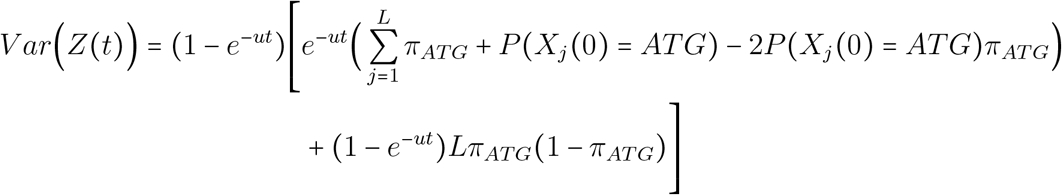

This expression can be further simplified by using the definitions for the quantities *Z_eq_*: = *Lπ_ATG_* and 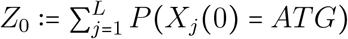, which results in the following form for the variance of ATGs:

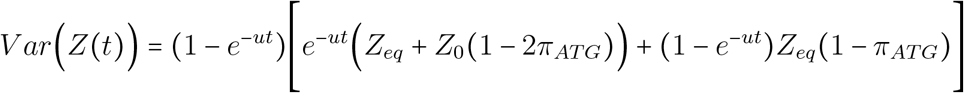

##### 5.3.2 Linear approximation for the variance of ATGs under F-81

In Section 3.4, we have shown that the variance at small time only depends on the parameter Ψ (see Equation 20):

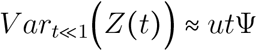

For this trait, it is straightforward to derive an analytical expression for Ψ and write it in terms of phenotypic quantities *Z_eq_* and *Z*_0_:

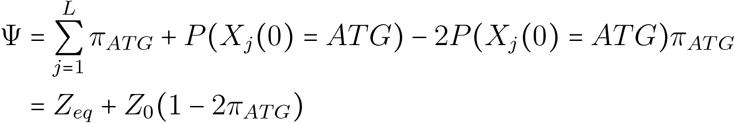

Note that assuming that the length of the sequence is known, *π_ATG_* can also be written in terms of phenotypic quantities, e.g. 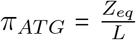. Given this expression for Ψ, we can write the linear approximation for the variance at short timescales as:

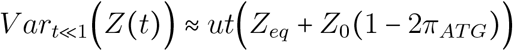

## References

[1] T. Bedford and D. L. Hartl, “Optimization of gene expression by natural selection,” Proc. Natl. Acad. Sci. U. S. A., vol. 106, no. 4, pp. 1133–1138, Jan. 2009, doi: 10.1073/pnas.0812009106.

[2] P. Khaitovich et al., “A Neutral Model of Transcriptome Evolution,” PLoS Biol., vol. 2, no. 5, May 2004, doi: 10.1371/journal.pbio.0020132.

[3] R. A. Studer et al., “Evolution of protein phosphorylation across 18 fungal species,” Science, vol. 354, no. 6309, pp. 229–232, 14 2016, doi: 10.1126/science.aaf2144.

[4] V. Mustonen, J. Kinney, C. G. Callan, and M. Lässig, “Energy-dependent fitness: a quantitative model for the evolution of yeast transcription factor binding sites,” Proc. Natl. Acad. Sci. U. S. A., vol. 105, no. 34, pp. 12376–12381, Aug. 2008, doi: 10.1073/pnas.0805909105.

[5] A. Nourmohammad, T. Held, and M. Lässig, “Universality and predictability in molecular quantitative genetics,” Curr. Opin. Genet. Dev., vol. 23, no. 6, pp. 684–693, Dec. 2013, doi: 10.1016/j.gde.2013.11.001.

[6] A. Nourmohammad, J. Rambeau, T. Held, V. Kovacova, J. Berg, and M. Lässig, “Adaptive Evolution of Gene Expression in Drosophila,” Cell Rep., vol. 20, no. 6, pp. 1385–1395, 08 2017, doi: 10.1016/j.celrep.2017.07.033.

[7] D. Brawand et al., “The evolution of gene expression levels in mammalian organs,” Nature, vol. 478, no. 7369, pp. 343–348, Oct. 2011, doi: 10.1038/nature10532.

[8] J. Chen et al., “A quantitative framework for characterizing the evolutionary history of mammalian gene expression,” Genome Res., vol. 29, no. 1, pp. 53–63, 2019, doi: 10.1101/gr.237636.118.

[9] J. G. Schraiber, Y. Mostovoy, T. Y. Hsu, and R. B. Brem, “Inferring Evolutionary Histories of Pathway Regulation from Transcriptional Profiling Data,” PLOS Comput. Biol., vol. 9, no. 10, p. e1003255, Oct. 2013, doi: 10.1371/journal.pcbi.1003255.

[10] T. F. Hansen, “Stabilizing Selection and the Comparative Analysis of Adaptation,” Evolution, vol. 51, no. 5, pp. 1341–1351, 1997, doi: 10.2307/2411186.

[11] R. Lande, “Natural Selection and Random Genetic Drift in Phenotypic Evolution,” Evolution, vol. 30, no. 2, pp. 314–334, Jun. 1976, doi: 10.2307/2407703.

[12] J. M. Beaulieu, D.-C. Jhwueng, C. Boettiger, and B. C. O’Meara, “Modeling stabilizing selection: expanding the ornstein–uhlenbeck model of adaptive evolution,” Evolution, vol. 66, no. 8, pp. 2369–2383, Aug. 2012, doi: 10.1111/j.1558-5646.2012.01619.x.

[13] M. A. Butler and A. A. King, “Phylogenetic Comparative Analysis: A Modeling Approach for Adaptive Evolution.,” Am. Nat., vol. 164, no. 6, pp. 683–695, Dec. 2004, doi: 10.1086/426002.

[14] J. C. Uyeda and J. E. and L. Harmon, bayou: Bayesian Fitting of Ornstein-Uhlenbeck Models to Phylogenies. 2018. [Online]. Available: https://cran.r-project.org/web/packages/bayou/index.html

[15] W.-C. Ho, Y. Ohya, and J. Zhang, “Testing the neutral hypothesis of phenotypic evolution,” Proc. Natl. Acad. Sci., vol. 114, no. 46, pp. 12219–12224, Nov. 2017, doi: 10.1073/pnas.1710351114.

[16] A. Whitehead and D. L. Crawford, “Neutral and adaptive variation in gene expression,” Proc. Natl. Acad. Sci., vol. 103, no. 14, pp. 5425–5430, Apr. 2006, doi: 10.1073/pnas.0507648103.

[17] J. Zhang, “Neutral Theory and Phenotypic Evolution,” Mol. Biol. Evol., vol. 35, no. 6, pp. 1327–1331, Jun. 2018, doi: 10.1093/molbev/msy065.

[18] J. Felsenstein, “Phylogenies and quantitative characters,” Annu. Rev. Ecol. Syst., vol. 19, no. 1, pp. 445–471, Nov. 1988, doi: 10.1146/annurev.es.19.110188.002305.

[19] M. Arenas, “Trends in substitution models of molecular evolution,” Front. Genet., vol. 6, Oct. 2015, doi: 10.3389/fgene.2015.00319.

[20] Z. Yang, Computational Molecular Evolution. Oxford, New York: Oxford University Press, 2006.

[21] R. A. Goldstein and D. D. Pollock, “Sequence entropy of folding and the absolute rate of amino acid substitutions,” Nat. Ecol. Evol., vol. 1, no. 12, pp. 1923–1930, Dec. 2017, doi: 10.1038/s41559-017-0338-9.

[22] Z. B. Zeng and C. C. Cockerham, “Mutation models and quantitative genetic variation.,” Genetics, vol. 133, no. 3, pp. 729–736, Mar. 1993.

[23] J. Felsenstein, “Maximum-likelihood estimation of evolutionary trees from continuous characters.,” Am. J. Hum. Genet., vol. 25, no. 5, pp. 471–492, Sep. 1973.

[24] J. Felsenstein, “Phylogenies and the Comparative Method,” Am. Nat., vol. 125, no. 1, pp. 1–15, 1985.

[25] R. Chakraborty and M. Nei, “Genetic differentiation of quantitative characters between populations or species: I. Mutation and random genetic drift,” Genet. Res., vol. 39, no. 3, pp. 303–314, Jun. 1982, doi: 10.1017/S0016672300020978.

[26] S. K. Davies, A. Leroi, A. Burt, J. G. Bundy, and C. F. Baer, “THE MUTATIONAL STRUCTURE OF METABOLISM IN CAENORHABDITIS ELEGANS,” Evol. Int. J. Org. Evol., vol. 70, no. 10, pp. 2239–2246, Oct. 2016, doi: 10.1111/evo.13020.

[27] J. F. C. Kingman, “A Simple Model for the Balance between Selection and Mutation,” J. Appl. Probab., vol. 15, no. 1, pp. 1–12, 1978, doi: 10.2307/3213231.

[28] A. Stoltzfus, “On the possibility of constructive neutral evolution,” J. Mol. Evol., vol. 49, no. 2, pp. 169–181, Aug. 1999, doi: 10.1007/pl00006540.

[29] M. Lynch, “The frailty of adaptive hypotheses for the origins of organismal complexity,” Proc. Natl. Acad. Sci. U. S. A., vol. 104 Suppl 1, pp. 8597–8604, May 2007, doi: 10.1073/pnas.0702207104.

[30] M. Lynch, “Phylogenetic divergence of cell biological features,” eLife, vol. 7, 21 2018, doi: 10.7554/eLife.34820.

[31] A. Nourmohammad, S. Schiffels, and M. Lässig, “Evolution of molecular phenotypes under stabilizing selection,” J. Stat. Mech. Theory Exp., vol. 2013, no. 01, p. P01012, Jan. 2013, doi: 10.1088/1742-5468/2013/01/P01012.

[32] B. Charlesworth, “Stabilizing selection, purifying selection, and mutational bias in finite populations,” Genetics, vol. 194, no. 4, pp. 955–971, Aug. 2013, doi: 10.1534/genetics.113.151555.

[33] L. M, “The Evolutionary Scaling of Cellular Traits Imposed by the Drift Barrier,” Proceedings of the National Academy of Sciences of the United States of America, May 12, 2020. https://pubmed.ncbi.nlm.nih.gov/32345718/ (accessed Jul. 03, 2020).

[34] A. Hodgins-Davis, D. P. Rice, and J. P. Townsend, “Gene Expression Evolves under a House-of-Cards Model of Stabilizing Selection,” Mol. Biol. Evol., vol. 32, no. 8, pp. 2130–2140, Aug. 2015, doi: 10.1093/molbev/msv094.

[35] A. Hodgins-Davis, F. Duveau, E. A. Walker, and P. J. Wittkopp, “Empirical measures of mutational effects define neutral models of regulatory evolution in Saccharomyces cerevisiae,” Proc. Natl. Acad. Sci. U. S. A., vol. 116, no. 42, pp. 21085–21093, 15 2019, doi: 10.1073/pnas.1902823116.

[36] B. Golding and J. Felsenstein, “A maximum likelihood approach to the detection of selection from a phylogeny,” J. Mol. Evol., vol. 31, no. 6, pp. 511–523, Dec. 1990, doi: 10.1007/BF02102078.

[37] J. Felsenstein, “Evolutionary trees from DNA sequences: A maximum likelihood approach,” J. Mol. Evol., vol. 17, no. 6, pp. 368–376, Nov. 1981, doi: 10.1007/BF01734359.

[38] W. J. Ewens, Mathematical Population Genetics 1: Theoretical Introduction, 2nd ed. New York: Springer-Verlag, 2004. doi: 10.1007/978-0-387-21822-9.

[39] M. Lynch, “The evolutionary scaling of cellular traits imposed by the drift barrier,” Proc. Natl. Acad. Sci., vol. 117, no. 19, pp. 10435–10444, May 2020, doi: 10.1073/pnas.2000446117.

[40] M. Kimura, “A simple method for estimating evolutionary rates of base substitutions through comparative studies of nucleotide sequences,” J. Mol. Evol., vol. 16, no. 2, pp. 111–120, Dec. 1980, doi: 10.1007/BF01731581.

[41] M. Lynch and W. G. Hill, “PHENOTYPIC EVOLUTION BY NEUTRAL MUTATION,” Evol. Int. J. Org. Evol., vol. 40, no. 5, pp. 915–935, Sep. 1986, doi: 10.1111/j.1558-5646.1986.tb00561.x.

[42] N. Cooper, G. H. Thomas, C. Venditti, A. Meade, and R. P. Freckleton, “A cautionary note on the use of Ornstein Uhlenbeck models in macroevolutionary studies,” Biol. J. Linn. Soc., vol. 118, no. 1, pp. 64–77, May 2016, doi: 10.1111/bij.12701.

[43] C. E. Cressler, M. A. Butler, and A. A. King, “Detecting Adaptive Evolution in Phylogenetic Comparative Analysis Using the Ornstein–Uhlenbeck Model,” Syst. Biol., vol. 64, no. 6, pp. 953–968, Nov. 2015, doi: 10.1093/sysbio/syv043.

[44] T. J. Meyer, J. L. Rosenkrantz, L. Carbone, and S. L. Chavez, “Endogenous Retroviruses: With Us and against Us,” Front. Chem., vol. 5, 2017, doi: 10.3389/fchem.2017.00023.

[45] P. Trávníček, M. Čertner, J. Ponert, Z. Chumová, J. Jersáková, and J. Suda, “Diversity in genome size and GC content shows adaptive potential in orchids and is closely linked to partial endoreplication, plant life-history traits and climatic conditions,” New Phytol., vol. 224, no. 4, pp. 1642–1656, Dec. 2019, doi: 10.1111/nph.15996.

[46] B. Paten et al., “Genome-wide nucleotide-level mammalian ancestor reconstruction,” Genome Res., vol. 18, no. 11, pp. 1829–1843, Nov. 2008, doi: 10.1101/gr.076521.108.

[47] K. A et al., “JASPAR 2018: Update of the Open-Access Database of Transcription Factor Binding Profiles and Its Web Framework,” Nucleic acids research, Jan. 04, 2018. https://pubmed.ncbi.nlm.nih.gov/29140473/ (accessed Jul. 11, 2020).

[48] B. Og and von H. Ph, “Selection of DNA Binding Sites by Regulatory Proteins. Statistical-mechanical Theory and Application to Operators and Promoters,” Journal of molecular biology, Feb. 20, 1987. https://pubmed.ncbi.nlm.nih.gov/3612791/ (accessed Jul. 11, 2020).

[49] W. W. Wasserman and A. Sandelin, “Applied bioinformatics for the identification of regulatory elements,” Nat. Rev. Genet., vol. 5, no. 4, pp. 276–287, Apr. 2004, doi: 10.1038/nrg1315.

[50] H.-D. A, D. F, W. Ea, and W. Pj, “Empirical Measures of Mutational Effects Define Neutral Models of Regulatory Evolution in Saccharomyces cerevisiae,” Proceedings of the National Academy of Sciences of the United States of America, Oct. 15, 2019. https://pubmed.ncbi.nlm.nih.gov/31570626/ (accessed Jul. 10, 2020).

[51] R. W. Lusk and M. B. Eisen, “Evolutionary Mirages: Selection on Binding Site Composition Creates the Illusion of Conserved Grammars in Drosophila Enhancers,” PLOS Genet., vol. 6, no. 1, p. e1000829, Jan. 2010, doi: 10.1371/journal.pgen.1000829.

[52] X. He, T. S. P. C. Duque, and S. Sinha, “Evolutionary Origins of Transcription Factor Binding Site Clusters,” Mol. Biol. Evol., vol. 29, no. 3, pp. 1059–1070, Mar. 2012, doi: 10.1093/molbev/msr277.

[53] B. A. Wilson, S. G. Foy, R. Neme, and J. Masel, “Young Genes are Highly Disordered as Predicted by the Preadaptation Hypothesis of De Novo Gene Birth,” Nat. Ecol. Evol., vol. 1, no. 6, p. 0146, Jun. 2017, doi: 10.1038/s41559-017-0146.

[54] W. Basile, O. Sachenkova, S. Light, and A. Elofsson, “High GC content causes orphan proteins to be intrinsically disordered,” PLoS Comput. Biol., vol. 13, no. 3, p. e1005375, 2017, doi: 10.1371/journal.pcbi.1005375.

[55] J. Prilusky et al., “FoldIndex: a simple tool to predict whether a given protein sequence is intrinsically unfolded,” Bioinforma. Oxf. Engl., vol. 21, no. 16, pp. 3435–3438, Aug. 2005, doi: 10.1093/bioinformatics/bti537.

[56] A. Lee, A. Nolan, J. Watson, and M. Tristem, “Identification of an ancient endogenous retrovirus, predating the divergence of the placental mammals,” Philos. Trans. R. Soc. B Biol. Sci., vol. 368, no. 1626, Sep. 2013, doi: 10.1098/rstb.2012.0503.

[57] J. R. Stone and G. A. Wray, “Rapid evolution of cis-regulatory sequences via local point mutations,” Mol. Biol. Evol., vol. 18, no. 9, pp. 1764–1770, Sep. 2001, doi: 10.1093/oxfordjournals.molbev.a003964.

[58] C. B, “Stabilizing Selection, Purifying Selection, and Mutational Bias in Finite Populations,” Genetics, Aug. 2013. https://pubmed.ncbi.nlm.nih.gov/23709636/ (accessed Jul. 10, 2020).

[59] Z. Yang and T. Zhu, “Bayesian selection of misspecified models is overconfident and may cause spurious posterior probabilities for phylogenetic trees,” Proc. Natl. Acad. Sci., vol. 115, no. 8, pp. 1854–1859, Feb. 2018, doi: 10.1073/pnas.1712673115.

[60] L. S. T. Ho and C. Ané, “Intrinsic inference difficulties for trait evolution with Ornstein-Uhlenbeck models,” Methods Ecol. Evol., vol. 5, no. 11, pp. 1133–1146, 2014, doi: 10.1111/2041-210X.12285.

[61] B. P. McEvoy and P. M. Visscher, “Genetics of human height,” Econ. Hum. Biol., vol. 7, no. 3, pp. 294–306, Dec. 2009, doi: 10.1016/j.ehb.2009.09.005.

[62] C. M. Jakobson and D. F. Jarosz, “Molecular Origins of Complex Heritability in Natural Genotype-to-Phenotype Relationships,” Cell Syst., vol. 8, no. 5, pp. 363–379.e3, 22 2019, doi: 10.1016/j.cels.2019.04.002.

[63] C. G. de Boer, E. D. Vaishnav, R. Sadeh, E. L. Abeyta, N. Friedman, and A. Regev, “Deciphering eukaryotic gene-regulatory logic with 100 million random promoters,” Nat. Biotechnol., vol. 38, no. 1, pp. 56–65, 2020, doi: 10.1038/s41587-019-0315-8.

[64] D. R. Kelley, Y. A. Reshef, M. Bileschi, D. Belanger, C. Y. McLean, and J. Snoek, “Sequential regulatory activity prediction across chromosomes with convolutional neural networks,” Genome Res., vol. 28, no. 5, pp. 739–750, 2018, doi: 10.1101/gr.227819.117.

[65] C. L. Strope, K. Abel, S. D. Scott, and E. N. Moriyama, “Biological sequence simulation for testing complex evolutionary hypotheses: indel-Seq-Gen version 2.0,” Mol. Biol. Evol., vol. 26, no. 11, pp. 2581–2593, Nov. 2009, doi: 10.1093/molbev/msp174.

[66] T. Zarin, C. N. Tsai, A. N. Nguyen Ba, and A. M. Moses, “Selection maintains signaling function of a highly diverged intrinsically disordered region,” Proc. Natl. Acad. Sci. U. S. A., vol. 114, no. 8, pp. E1450–E1459, 21 2017, doi: 10.1073/pnas.1614787114.

[67] T. Zarin, B. Strome, A. N. Nguyen Ba, S. Alberti, J. D. Forman-Kay, and A. M. Moses, “Proteome-wide signatures of function in highly diverged intrinsically disordered regions,” eLife, vol. 8, Jul. 2019, doi: 10.7554/eLife.46883.

[68] S. Nakagawa and M. U. Takahashi, “gEVE: a genome-based endogenous viral element database provides comprehensive viral protein-coding sequences in mammalian genomes,” Database J. Biol. Databases Curation, vol. 2016, 2016, doi: 10.1093/database/baw087.

[69] A. Yates et al., “The Ensembl REST API: Ensembl Data for Any Language,” Bioinformatics, vol. 31, no. 1, pp. 143–145, Jan. 2015, doi: 10.1093/bioinformatics/btu613.

[70] K. P. Schliep, “phangorn: phylogenetic analysis in R,” Bioinformatics, vol. 27, no. 4, pp. 592–593, Feb. 2011, doi: 10.1093/bioinformatics/btq706.

[71] A. Moses and S. Sinha, “Regulatory Motif Analysis,” in Bioinformatics: Tools and Applications, 2009, pp. 137–163. doi: 10.1007/978-0-387-92738-1_7.

[72] D. R. Zerbino et al., “Ensembl 2018,” Nucleic Acids Res., vol. 46, no. D1, pp. D754–D761, 04 2018, doi: 10.1093/nar/gkx1098.

[73] Z. Yang, “PAML 4: phylogenetic analysis by maximum likelihood,” Mol. Biol. Evol., vol. 24, no. 8, pp. 1586–1591, Aug. 2007, doi: 10.1093/molbev/msm088.

[74] E. Paradis and K. Schliep, “ape 5.0: an environment for modern phylogenetics and evolutionary analyses in R,” Bioinforma. Oxf. Engl., vol. 35, no. 3, pp. 526–528, 01 2019, doi: 10.1093/bioinformatics/bty633.

[75] P. Virtanen et al., “SciPy 1.0: fundamental algorithms for scientific computing in Python,” Nat. Methods, vol. 17, no. 3, pp. 261–272, Mar. 2020, doi: 10.1038/s41592-019-0686-2.

[76] M. W. Pennell et al., “geiger v2.0: an expanded suite of methods for fitting macroevolutionary models to phylogenetic trees,” Bioinformatics, vol. 30, no. 15, pp. 2216–2218, Aug. 2014, doi: 10.1093/bioinformatics/btu181.

## References

[1] Marguerite A. Butler and Aaron A. King. Phylogenetic comparative analysis: a modeling approach for adaptive evolution. American Naturalist, 164:683–695, 2004.

[2] Natalie Cooper, Gavin H. Thomas, Chris Venditti, Andrew Meade, and Rob P. Freckleton. A cautionary note on the use of Ornstein Uhlenbeck models in macroevolutionary studies. Biological Journal of the Linnean Society, 118(1), 2016.

[3] Will Cornwell and Shinichi Nakagawa. Phylogenetic comparative methods, 2017.

[4] Warren J. Ewens. Mathematical Population Genetics: I. Theoretical Introduction. 2004.

[5] Joseph Felsenstein. Evolutionary trees from DNA sequences: A maximum likelihood approach. Journal of Molecular Evolution, 17(6), 1981.

[6] Joseph Felsenstein. Theoretical evolutionary genetics. 2019.

[7] Luke J. Harmon. Phylogenetic Comparative Methods: Learning from Trees. 2018.

[8] Torsten Held, Armita Nourmohammad, and Michael Lässig. Adaptive evolution of molecular phenotypes. Journal of Statistical Mechanics: Theory and Experiment, 2014(9), 2014.

[9] T H Jukes and C R Cantor. Evolution of protein molecules BT - Mammalian protein metabolism. In Mammalian protein metabolism, volume III. 1969.

[10] Armita Nourmohammad, Joachim Rambeau, Torsten Held, Viera Kovacova, Johannes Berg, and Michael Lässig. Adaptive Evolution of Gene Expression in Drosophila. Cell Reports, 20(6), 2017.

[11] Armita Nourmohammad, Stephan Schiffels, and Michael Lässig. Evolution of molecular phenotypes under stabilizing selection. Journal of Statistical Mechanics: Theory and Experiment, 2013(1), 2013.

[12] Matthew W. Pennell, Jonathan M. Eastman, Graham J. Slater, Joseph W. Brown, Josef C. Uyeda, Richard G. Fitzjohn, Michael E. Alfaro, and Luke J. Harmon. Geiger v2.0: An expanded suite of methods for fitting macroevolutionary models to phylogenetic trees. Bioinformatics, 30(15), 2014.

[13] Lam Si Tung Ho and Cecile Ane. A linear-time algorithm for gaussian and nongaussian trait evolution models. Systematic Biology, 63(3), 2014.

[14] Ziheng Yang. Computational Molecular Evolution. 2010.

